# Spacing effect improves generalization in biological and artificial systems

**DOI:** 10.64898/2025.12.18.695340

**Authors:** Guanglong Sun, Ning Huang, Hongwei Yan, Jun Zhou, Qian Li, Bo Lei, Yi Zhong, Liyuan Wang

## Abstract

Generalization is a fundamental criterion for evaluating learning effectiveness, a domain where biological intelligence excels yet artificial intelligence continues to face challenges. In biological learning and memory, the well-documented *spacing effect* shows that appropriately spaced intervals between learning trials can significantly improve behavioral performance. While multiple theories have been proposed to explain its underlying mechanisms, one compelling hypothesis is that spaced training promotes integration of input and innate variations, thereby enhancing generalization to novel but related scenarios. Here we examine this hypothesis by introducing a bio-inspired spacing effect into artificial neural networks, integrating input and innate variations across spaced intervals at the neuronal, synaptic, and network levels. These spaced ensemble strategies yield significant performance gains across various benchmark datasets and network architectures. Biological experiments on *Drosophila* further validate the complementary effect of appropriate variations and spaced intervals in improving generalization, which together reveal a convergent computational principle shared by biological learning and machine learning.

## INTRODUCTION

The ability to generalize previously learned knowledge to novel but related scenarios is a hallmark of intelligent behavior. For example, a person who has learned to recognize another’s voice can identify it again despite variations in background noise, pitch, or speaking speed. This underlies adaptive decision-making in everyday life and represents an almost innate advantage of biological intelligence (BI). In comparison, generalization from training data to similar test data is a fundamental objective of artificial intelligence (AI), especially for modern artificial neural networks (ANNs)-based systems^1,2^. Understanding the mechanisms that support robust generalization in BI offers a promising path toward improving generalization in AI. Conversely, implementing bio-inspired generalization strategies in ANNs allows for reverse-engineering and computational validation of candidate biological hypotheses. This interdisciplinary research paradigm, known as NeuroAI^1–4^, holds promise for uncovering their convergent computational principles.

One well-documented phenomenon in biological learning and memory is the *spacing effect* ^5^: spaced training with scattered repetitions over a certain period of time is significantly more effective than massed training with focused repetitions. This effect has been reported across species, from invertebrates to humans^6–13^, and across a wide range of representative learning tasks^14–19^. Several cognitive theories have been proposed to account for this effect^20^. The *deficient-processing* theory^7,21–23^ suggests that massed repetitions allow insufficient time for essential biological processes such as protein synthesis and synaptic plasticity, thereby impairing knowledge encoding. The *study-phase retrieval* theory^24,25^ posits that spaced trials facilitate retrieval and strengthening of earlier memory traces, thereby enhancing knowledge retention. The *encoding variability* theory^26–29^ suggests that spacing can introduce natural variations in inputs or internal states, producing a more diverse set of memory traces. Importantly, these accounts are not mutually exclusive and may be complementary. Among them, encoding variability emphasizes how structured variations accumulated across repetitions can support robust responses beyond the specific training instance, providing a comparably explicit handle on *generalization*.

A parallel mechanism for improving generalization has emerged in machine learning, known as ensemble learning (EL)^2,4,30–32^, which aggregates multiple models that capture diverse aspects of the data distribution. Similar to the temporally evolving nature of biological learning, temporal ensemble strategies^33,34^ leverage variability accumulated across training steps to construct more robust and generalizable representations (Fig. 1A,B), which include three representative implementations: (1) dropout^35^ introduces stochasticity at the *neuronal level* by randomly deactivating units during training, promoting the formation of diverse activation patterns; (2) weight averaging (WA)^36^, often implemented via exponential moving average (EMA)^37^, aggregates parameter snapshots throughout network training, which captures variability at the *synaptic level*; (3) knowledge distillation (KD)^38^, especially online KD^39^ and self KD^40^, transfers soft output distributions from temporally more advanced teacher models to student models, integrating variability at the *network level*.

**Figure 1.**
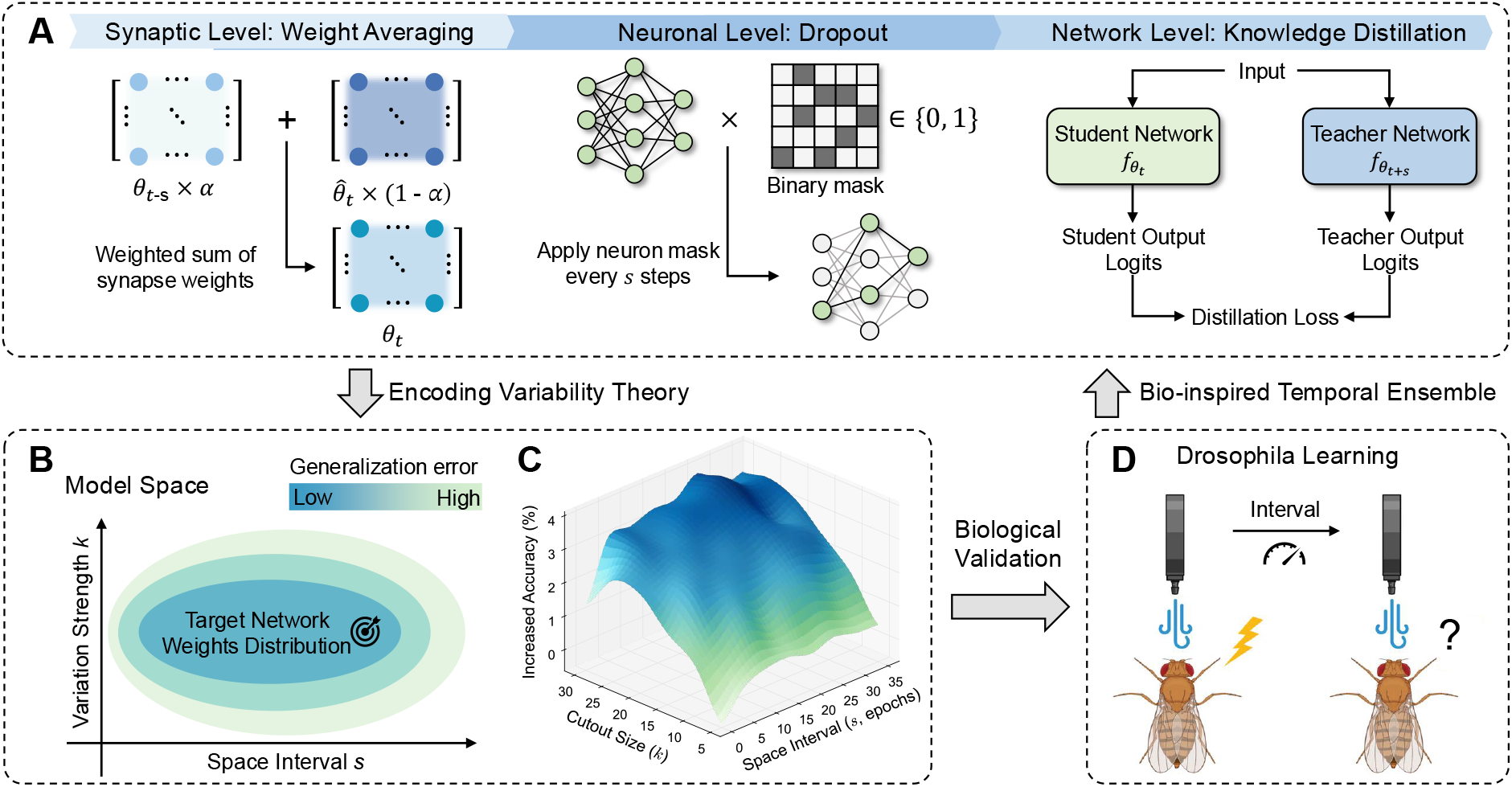
Overview of spacing effect in biological and artificial systems. (**A**) Schematic illustration of temporal ensemble strategies at the neuronal, synaptic, and network levels in ANNs, capturing the temporally evolving nature of biological learning. (**B**) Two key factors, variation strength and spaced interval, jointly modulate generalization performance in an independent and complementary manner. (**C**) Joint analysis of variation strength and spaced interval with cutout-based augmentation on CIFAR-10 dataset reveals an inverted U-shaped performance profile with respect to both factors. All results are averaged over three runs with different random seeds. (**D**) *Drosophila* learning with olfactory-conditioning experiments.

Based on a unified formulation of temporal ensemble strategies, we revisit and computationally validate the encoding variability theory that may underlie the spacing effect in biological systems. Specifically, we incorporate bio-inspired spacing effects into ANNs at the neuronal, synaptic, and network levels (corresponding to dropout^35^, WA^36^, and KD^38^, respectively) by introducing controllable input and innate variations across spaced training intervals. Across a variety of benchmark datasets and network architectures, we observe a consistent inverted Ushaped relationship of the training-to-test generalization with both the magnitude of variations and the duration of spaced intervals. Strikingly, either appropriate variation strength under fixed spaced interval or appropriate spaced interval under fixed variation strength leads to considerable improvements in test performance. These findings point to a critical balance between structured variability and temporal spacing for enhanced generalization.

Guided by these computational insights, we further design olfactory-conditioning experiments with *Drosophila* to assess whether biological generalization similarly benefits from this balance. We find that either increasing the spacing between repetitions or introducing variability in training cues significantly improves test performance in scenarios that differ slightly from training, suggesting that biological generalization also benefits from the structured integration of variations over a certain period of time. These findings demonstrate a shared computational principle underlying biological and artificial systems: structured exposure to appropriate variations promotes generalization from previous experiences. This principle opens up promising directions for designing biologically grounded training paradigms in ANNs, while offering a computational lens to understand naturally evolved “algorithms” in biological brains.

## RESULTS

### Spaced temporal ensemble of encoding variability in biological and artificial systems

The encoding variability theory in BI posits that generalization is enhanced by appropriately spaced repetitions that introduce diverse representations, either through external inputs or internal neural states^7,26,27^. A parallel mechanism exists in AI, where EL improves generalization by aggregating appropriately diversified model variants that capture complementary aspects of the data distribution. Such diversity can arise from input variations introduced through external data augmentation^41^ or from innate variations introduced throughout training. The innate variations are typically instantiated via temporal ensemble strategies such as dropout^35^, WA^36^, and KD^38^, which operate at the neuronal, synaptic, and network levels, respectively.

In both biological and artificial systems, the effectiveness of variability in improving generalization critically depends on its magnitude and the temporal spacing over which it is integrated. This observation leads us to hypothesize a shared computational principle: structured exposure to appropriate variations over a certain period of time enhances generalization from training to test. To evaluate this, we systematically investigate how input and innate variations, modulated by controllable spaced intervals, affect generalization performance in ANNs, and further examine whether analogous effects occur in biological learning and memory through behavioral experiments in *Drosophila*.

### Spacing effect with input and innate variations enhances generalization in ANNs

We begin by investigating the spacing effect in ANNs using externally imposed **input variations**. Specifically, we adopt the cutout augmentation strategy^42^, which simulates environmental perturbations by randomly masking square patches in input images. This strategy allows independent manipulation of two key factors: the *variation strength* defined by the cutout size (Supplementary Fig. S1A), and the *spaced interval* defined by the frequency at which the cutout size is changed during training (Supplementary Fig. S1B). In ANNs, we use “spaced interval” to denote an interval in training steps or epochs between variability updates (e.g., changing cutout size), rather than a physical delay between learning events as in behavioral experiments. For all ANNs experiments, we follow the standard image-classification protocol: models are trained on the official training set and evaluated on a disjoint held-out test set (*no overlapping images*) drawn from the same underlying distribution. Accordingly, “generalization” here refers to training-to-test generalization within each benchmark dataset. To further contextualize this setting, we quantify similarity between training and test sets with the Wasserstein distance metric (Supplementary Fig. S2). See Discussion for an extensive comparison of the “generalization” concept between biological learning and machine learning.

Using a 4-layer convolutional neural network (CNN) trained on CIFAR-10 dataset^43^, we investigate how test performance varies with cutout size under a fixed augmentation schedule. We observe a clear inverted U-shaped trend: moderate masking enhances generalization from training to test, whereas overly large occlusions impair the performance (Supplementary Fig. S1C). Next, we introduce temporal structure by periodically altering the cutout size across training epochs. Such a spaced version of cutout augmentation further improves test performance and again yields an inverted U-shaped trend, now with respect to the spaced interval (Supplementary Fig. S1D). Notably, the effects of variation strength and spaced interval appear largely independent and additive, suggesting that both factors contribute complementary benefits to generalization. When combined, they jointly promote more robust representations and improved test performance (Fig. 1C).

Recent theoretical work^44^ has demonstrated that representative temporal ensemble strategies^33,34^, including dropout^35^, EMA^37^, online KD^38^, and self KD^40^, are fundamentally equivalent in improving generalization, and all amount to integrating properly diversified model variants throughout training (Fig. 1B). Building upon this foundation, we introduce a unified formulation that explicitly organizes temporal ensemble strategies under the common principle of spaced ensembling, governed by the *spaced interval s* between ensemble updates and the *variation strength k* applied at each update:

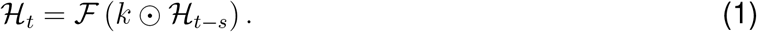

ℋ _*t*_ denotes the internal state at the current training step *t*, which may refer to neuron activations *h*_*t*_, network parameters *θ*_*t*_, or model outputs 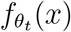 given inputs *x*. The operator ⊙ represents element-wise modulation. ℱ is an ensembling function adapted to each strategy. Throughout the paper, *t* indexes the training steps or epochs, and *s* denotes the interval along the training trajectory between ensemble updates, rather than a physical inter-stimulus time. Here, *k* serves as a possibly high-dimensional *variation operator* that may encode both the *magnitude* and *structure* of variations (e.g., cutout masks, dropout masks, or teacher-student pairing). In our controlled sweeps we summarize this operator by a scalar “variation strength” for clarity. With this unified formulation, we empirically investigate the bio-inspired spacing effect in ANNs across three levels of innate variations.

#### Neuronal Level: Dropout

Dropout^35^ introduces innate variations by randomly deactivating a subset of neurons throughout training. Here, the *variation strength* corresponds to the dropout rate (i.e., the probability of deactivation), while the *spaced interval* is inherently fixed, as dropout is often applied at every training step. To explore the impact of temporal structure, we employ a periodic schedule on the dropout rate as the spaced version of dropout (Fig. 2A, Supplementary Algorithm 1), akin to the idea of implementing the spacing effect with cutout augmentation. Here, the spaced interval *s* is implemented as the period in training steps or epochs over which the dropout probability is modulated. The training remains continuous, without inserting physical gaps between mini-batches.

**Figure 2.**
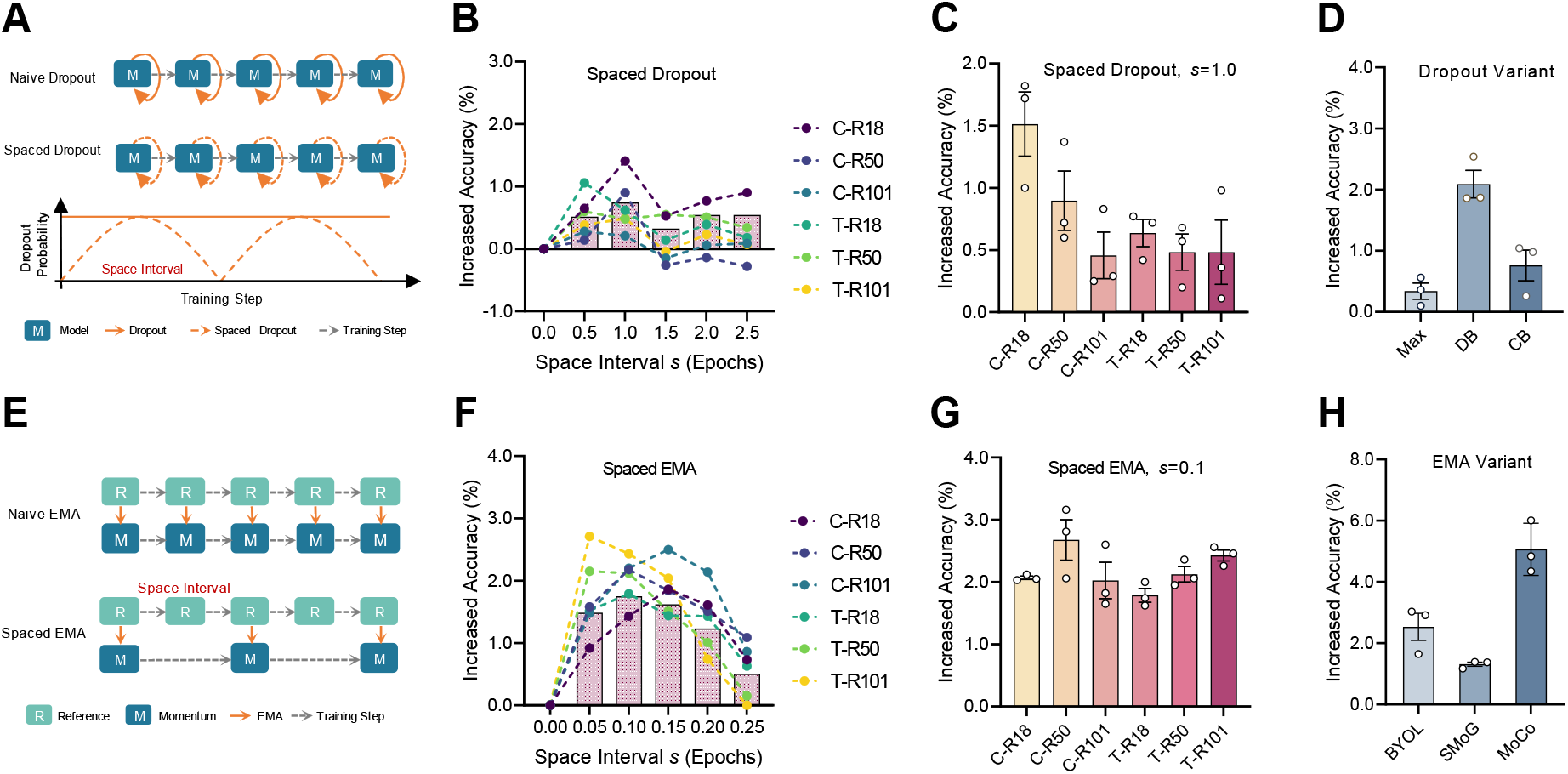
Spacing effect enhances generalization through structured dropout and WA. (**A**) Spaced version of dropout, where the dropout probability varies periodically to introduce structured neuronal variability during training. (**B**) Performance gains under different structured dropout probabilities exhibit an inverted U-shaped trend, indicating optimal performance at intermediate variation strengths. The spaced interval *s* = 0 denotes the baseline performance without spacing effect modification. (**C**) Performance gains of spaced dropout across different network architectures and benchmark datasets. (**D**) Spaced version of dropout improves performance in advanced dropout variants, including MaxDropout^45^, DropBlock^46^, and Checkerboard^47^. (**E**) Spaced version of EMA, where momentum updates occur at fixed intervals to control synaptic variability. (**F**) Performance gains under different momentum spacing intervals also exhibit an inverted U-shaped trend, highlighting the importance of optimal temporal spacing. The spaced interval *s* = 0 denotes the baseline performance without spacing effect modification. (**G**) Performance gains of spaced EMA across different network architectures and benchmark datasets. (**H**) Spaced version of EMA improves performance in advanced EMA variants with self-supervised learning, including BYOL^48^, SMoG^49^, and MoCo^50^. C, CIFAR-100. T, Tiny-ImageNet. R18, ResNet-18. R50, ResNet-50. R101, ResNet-101. All results are averaged over three runs with different random seeds. Data are presented as mean ± SEM. See Supplementary Table S1 for the original results of baselines and spaced variants.

We observe a clear inverted U-shaped relationship between test performance and dropout rate (Fig. 2B): mild deactivation promotes diverse activations and therefore improves generalization, whereas excessive sparsity of neuronal activations impairs the model’s learning capacity. Importantly, this inverted U-shaped trend persists when jointly manipulating the spaced interval *s* and the dropout rate as the variation strength *k* (Supplementary Fig. S3A), mirroring the spac-ing effect observed in input variations. The spaced version of dropout consistently outperforms standard dropout (Fig. 2C) across multiple benchmark datasets (CIFAR-100 and Tiny-ImageNet) and network architectures (ResNet-18, -50, and -101). The benefits generalize across advanced dropout variants including DropBlock^46^, MaxDropout^45^, and Checkerboard^47^ (Fig. 2D). These results underscore the robustness of temporally structured neuronal variations in improving generalization.

#### Synaptic Level: Weight Averaging

WA^36^ such as EMA^37^ enhances generalization by aggregating temporally spaced parameter snapshots throughout training. Here, the *spaced interval* is implicitly determined by the EMA momentum coefficient, which controls the effective temporal distance between aggregated snapshots (Fig. 2E, Supplementary Algorithm 2). In this context, the temporal distance refers to separation along the training steps or epochs between aggregated parameter states, rather than elapsed physical time between learning events. The *variation strength* corresponds to the magnitude of parameter updates, which is not directly tunable in standard EMA.

By modulating the momentum value, we again observe a clear inverted U-shaped relationship between test performance and spaced interval (Fig. 2F): overly short intervals produce highly similar snapshots with limited diversity, while overly long intervals lead to incoherent aggregation that compromises performance. As shown in Supplementary Fig. S3B, different momentum values (serving here as the variation strength *k*) substantially affect the location of performance peak: lower momentum accelerates diversity but impairs stability, whereas higher momentum suppresses useful variations. The spaced version of EMA achieves consistent performance gains across multiple benchmark datasets and network architectures (Fig. 2G), and extends to more advanced EMA variants such as BYOL^48^, SMoG^49^ and MoCo^50^ (Fig. 2H). These results validate the benefits of temporally spaced integration of encoding variability at the synaptic level.

#### Network Level: Knowledge Distillation

KD^38^ improves generalization by training student models with the soft outputs of more advanced teacher models. We investigate two representative temporal variants: online KD^39^, where the teacher is asynchronously updated throughout training; and self KD^40^, where the student periodically supervises itself using earlier snapshots. In both cases, the *spaced interval* corresponds to the number of epochs between teacher updates, while the *variation strength* naturally arises from the divergence between teacher and student over time. Therefore, the spaced interval in KD is implemented as an update frequency for teacher–student alignment, rather than elapsed physical time between learning events.

For online KD (Fig. 3A, Supplementary Algorithm 3), we observe a consistent inverted Ushaped relationship between test performance and spaced interval (Fig. 3B): overly short intervals offer limited variations, while overly long intervals lead to teacher-student mismatch and degraded performance. A similar trend emerges in self KD (Fig. 3E–H, Supplementary Algorithm 4), where periodic re-alignment with earlier snapshots again exhibits an inverted U-shaped performance profile (Fig. 3F). Further, Supplementary Fig. S3C confirms that adaptation of variation strength (e.g., via data shuffling) affects the optimal spaced interval, with stronger variations favoring shorter intervals. The spaced versions of online KD (Fig. 3C,D) and self KD (Fig. 3G,H) yield consistent performance gains across multiple benchmark datasets, network architectures, and advanced variants (e.g., SHAKE^51^, CTKD^52^, and LSKD^53^ for online KD; and DLB^54^, PSKD^55^ and TSB^56^ for self KD), suggesting the effectiveness of bio-inspired spacing effect at the network level.

**Figure 3.**
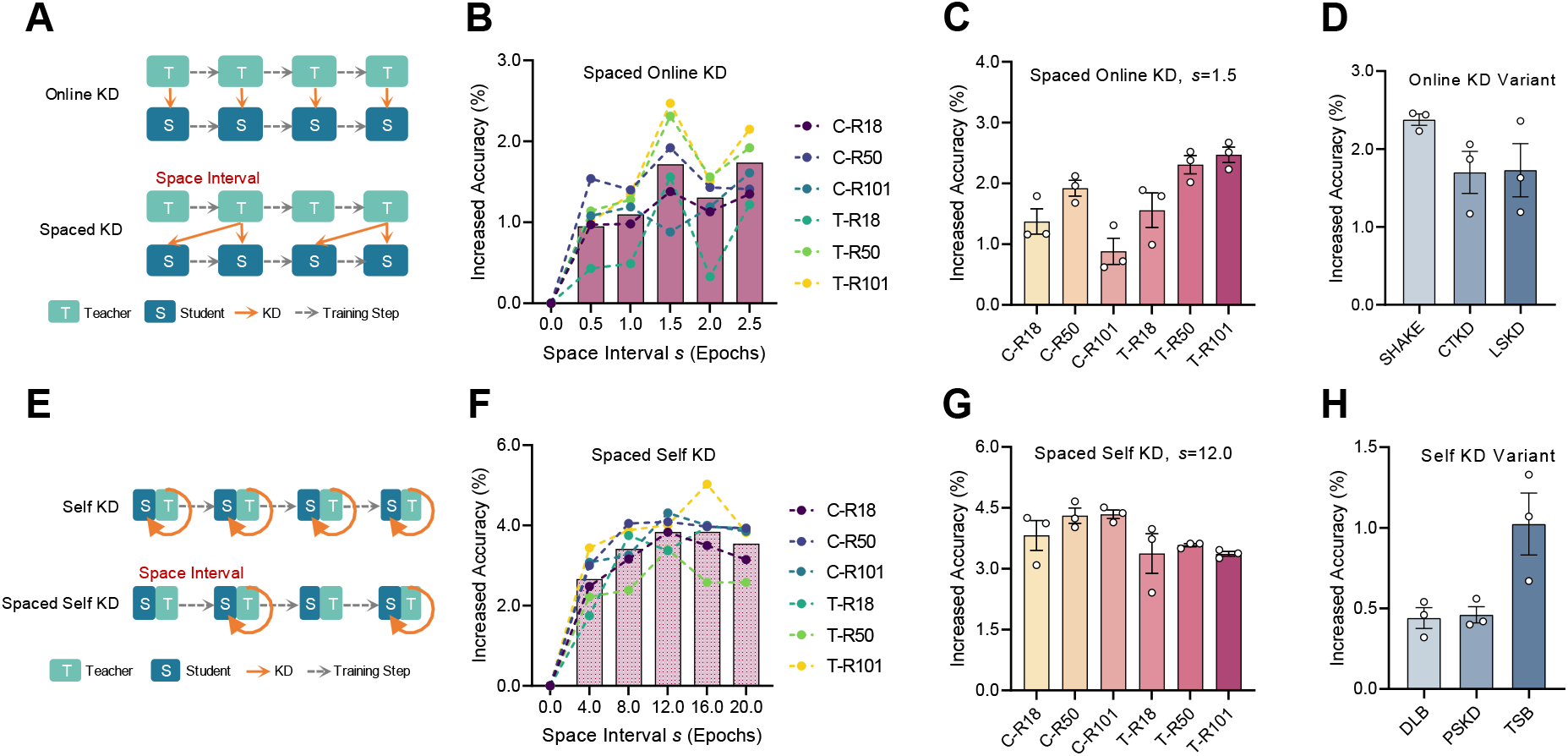
Spacing effect enhances generalization in online KD and self KD. (**A**) Spaced version of online KD, where the teacher model is updated at fixed intervals to induce innate variations. (**B**) Performance gains of spaced online KD exhibit an inverted U-shaped trend with respect to spaced interval. The spaced interval *s* = 0 denotes the baseline performance without spacing effect modification. (**C**) Performance gains of spaced online KD across different network architectures and benchmark datasets. (**D**) Spaced version of online KD improves performance in advanced online KD variants, including SHAKE^51^, CTKD^52^, and LSKD^53^. (**E**) Spaced version of self KD, where the deepest layers (teacher) periodically supervise shallower layers (students) with fixed intervals. (**F**) Performance gains of spaced self KD exhibit an inverted U-shaped trend with respect to spaced interval. The spaced interval *s* = 0 denotes the baseline performance without spacing effect modification. (**G**) Performance gains of spaced self KD across different network architectures and benchmark datasets. (**H**) Spaced version of self KD improves performance in advanced self KD variants, including DLB^54^, PSKD^55^ and TSB^56^. C, CIFAR-100. T, Tiny-ImageNet. R18, ResNet-18. R50, ResNet-50. R101, ResNet-101. All results are averaged over three runs with different random seeds. Data are presented as mean ± SEM. See Supplementary Table S1 for the original results of baselines and spaced variants.

Together, these results demonstrate that appropriately spaced exposure to either input or innate variations, whether introduced through data augmentation^42^, dropout^35^, WA^36^, online KD^38^, and self KD^40^, consistently enhances generalization from training to test in ANNs. Moreover, Supplementary Fig. S3 highlights a consistent inverted U-shaped relationship between variation strength *k* and spaced interval *s* across all methods, reinforcing the unified role of spacing-induced encoding variability from a machine learning perspective.

Additionally, we examine whether spacing effects at different levels are complementary by evaluating combinations of the spaced variants of dropout, EMA, and KD. We observe cumulative performance gains when combining spaced dropout and spaced EMA relative to using either strategy alone. This suggests that imposing spacing concurrently at the neuronal and synaptic levels introduces partially non-overlapping sources of variation that further improve generalization. However, we note that the spaced KD condition does not yield further additive benefits when combined with these other spacing effects (Supplementary Fig. S4). To test whether the performance gains arise from the periodic structure of spacing, we compare the periodic spaced schedules against matched random schedules in which the same variability updates are applied at irregular, shuffled intervals while preserving the same average update frequency (Supplementary Fig. S5). The periodic spaced schedules consistently outperform the random counterparts across temporal ensemble strategies, benchmark datasets, and network architectures. These results suggest the periodic structure of spacing plays an important role in improving generaliza-tion, consistent with the precise timing requirements observed in biological spacing effects^5,7,20^. Finally, we characterize how spacing influences the training trajectory. Across temporal ensemble strategies, spaced training does not noticeably increase early-stage learning speed, but it consistently raises the converged performance ceiling (Supplementary Fig. S6), leading to higher final test accuracy. These improvements remain consistent under different magnitudes of parameter perturbations by adding Gaussian noise (Supplementary Fig. S7), suggesting the robustness of our parameterized solution in adapting to changes.

### Spaced training and trial-to-trial variation enhance generalization in *Drosophila*

To investigate whether the unified role of spacing-induced encoding variability in ANNs also applies to biological systems, we train the *Drosophila* fruit flies using a classical olfactory aversive conditioning paradigm and assess both conditioned memory and its generalization performance^57–59^ (Supplementary Fig. S8A,B). In a forward training (FT) trial, flies are exposed to the odor 3-OCT paired with electric shocks (conditioned stimulus plus, CS+), followed by the odor EA without shock (conditioned stimulus minus, CS-). Both odor exposure only (OEO) trial and backward training (BT) trial serve as controls for evaluating associative memory performance. After training, conditioned and generalized memory are tested by presenting either the CS+ odor (3-OCT) or a structurally similar odor (1-OCT), respectively, against a novel odor (MCH). Our experiments validate that memory formation is associative and that flies can generalize to the similar odor as early as 3 min after training (Supplementary Fig. S8C-E).

First, we examine the effect of temporal spacing on memory generalization. We train flies with five repeated trials separated by different inter-trial intervals (ITIs), ranging from massed training (45-s ITI) to spaced training (5-, 15-, 30-, or 60-min ITI; Fig. 4A). At 3 min after training, all groups exhibit robust conditioned and generalized memory, with no significant differences across variable ITI conditions (Fig. 4B,C). The generalization ratio also has no significant difference among groups (Fig. 4D). In contrast, at 24 hours after training, both conditioned and generalized memory are significantly influenced by ITI variation. Specifically, flies subjected to spaced training with 15-min ITIs show significantly enhanced conditioned and generalized memory compared to the massed training group (Fig. 4E,F), resulting in an inverted U-shaped pattern across ITI groups (Fig. 4G). These results underscore the importance of spaced intervals in promoting memory generalization.

**Figure 4.**
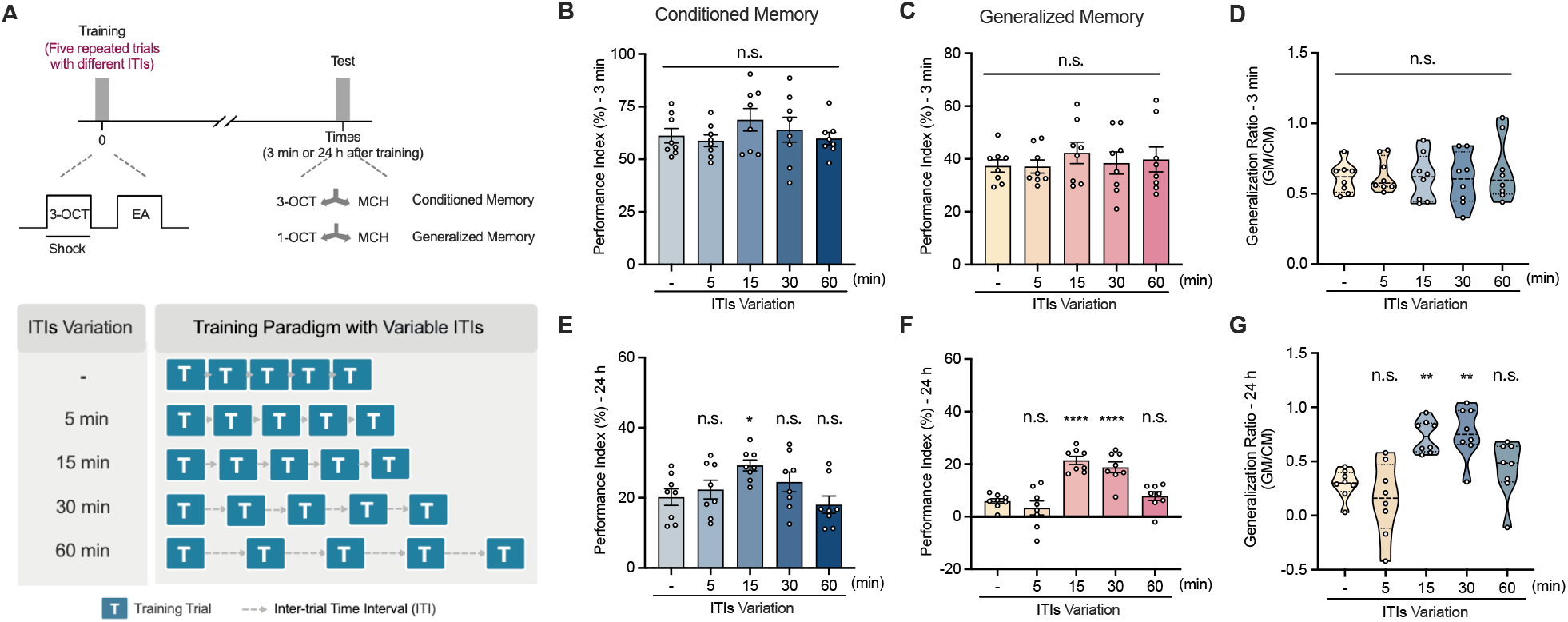
Spaced training enhances memory generalization in *Drosophila*. (**A**) Experimental paradigm for aversive olfactory conditioning with variable inter-trial intervals (ITI). The massed training group (-) receives five trials with a 45-s ITI, while the spaced training groups (5, 15, 30, 60 min) receive trials with increasing ITIs. Conditioned and generalized memory are tested by presenting the CS+ odor (3-OCT) or a structurally similar odor (1-OCT), respectively, against a novel odor (MCH). (**B, C**) Conditioned memory (**B**, n=8) and generalized memory (**C**, n=8) at 3 min show no significant difference across ITI groups. (**D**) The generalization ratio shows no significant difference across ITI groups at 3 min. n=8. (**E, F**) Conditioned memory (**E**, n=8) and generalized memory (**F**, n=8) at 24 h exhibit an inverted U-shaped pattern across ITI groups. (**G**) The generalization ratio is significantly improved in the 15- and 30-min ITI groups at 24 h. n=8. Data are presented as mean ± SEM. Statistical significance is determined by one-way ANOVA with Dunnett’s multiple comparisons test. *p≤0.05, **p≤0.01, ****p≤0.0001, n.s., not significant.

Next, we investigate the impact of encoding variability, inspired by our findings in machine learning. We introduce trial-to-trial variation during massed training by systematically varying the odor delivery flow rate (0.5, 1, or 1.5 K ml/min) across five trials (Fig. 5A). This manipulation does not impair conditioned memory, as all variable-training groups performed comparably to the constant-flow control group (Fig. 5B). Strikingly, introducing variability significantly enhances generalized memory (Fig. 5C) and increases the generalization ratio (Fig. 5D). This benefit is also persistent, as the enhancement in generalized memory is still evident 24 hours after training (Fig. 5E,F). However, varying shock intensities across trials does not affect generalization under these conditions (Supplementary Fig. S9), indicating that variability in the conditioned sensory input (CS), rather than variability in the reinforcement strength (US), is the critical factor driving improved generalization.

**Figure 5.**
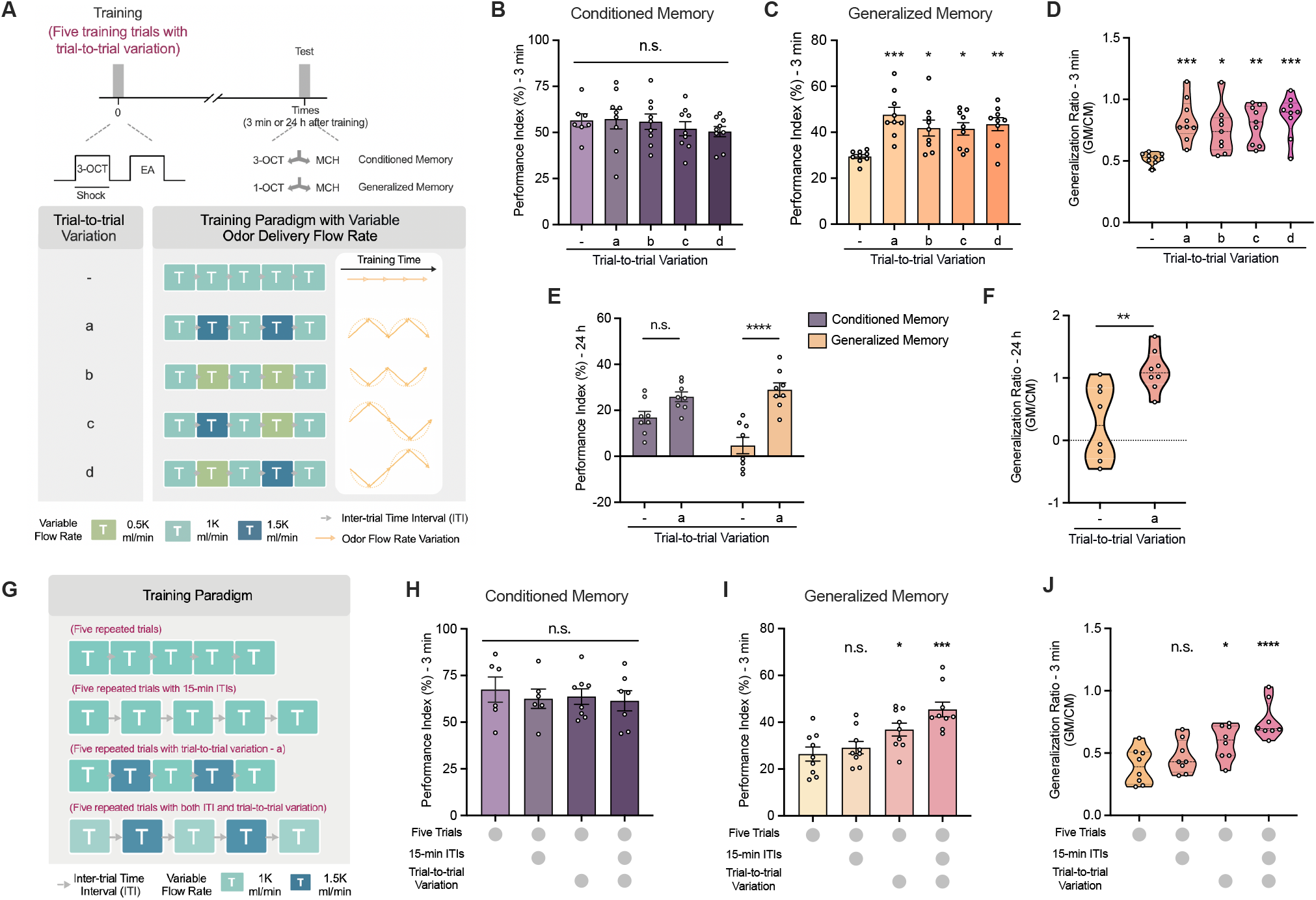
Trial-to-trial variation enhances memory generalization in *Drosophila*. (**A**) Experimental paradigm for massed training with variable sensory input. The control group (-) receives five trials with a constant odor delivery flow rate, while the experimental groups (a-d) receive trials with varying flow rates to introduce encoding variability. (**B**) Conditioned memory shows no significant difference between control and experimental groups (a-d) at 3 min. n=7-9. (**C**) All experimental groups (a-d) exhibit enhanced generalized memory at 3 min. n=9. (**D**) The generalization ratio is significantly increased in experimental groups (a-d) at 3 min. n=9. (**E**) Generalized memory remains enhanced in the experimental group (a) compared to the control group (-) at 24 h. n=8. (**F**) The generalization ratio remains elevated in the experimental group (a) at 24 h. n=8. (**G**) Experimental paradigm for evaluating the interaction between temporal spacing and trial-to-trial variation: five repeated trials (massed), five trials with trial-to-trial variation, five trials with 15-min ITIs (spaced), and five trials combining both 15-min ITIs and trial-to-trial variation. (**H**) Conditioned memory shows no significant difference across all groups at 3 min. n=5. (**I, J**) Generalized memory (**I**) and generalization ratio (**J**) at 3 min. Data are presented as mean ± SEM. Statistical significance is determined by one-way ANOVA with Dunnett’s multiple comparisons test. *p≤0.05, **p≤0.01, ***p≤0.001, ****p≤0.0001, n.s., not significant.

To further explore the interplay between encoding variability and classical biological spaced learning, we examine whether trial-to-trial variation interacts with ITIs (Fig. 5G). Beyond the benefits of encoding variability observed during massed training (45-s ITIs), we find that extending the ITIs to 15 min also significantly enhances memory generalization. Remarkably, combining both 15-min ITIs and trial-to-trial variation yields a synergistic improvement in both generalized memory and the generalization ratio (Fig. 5I, J), surpassing the effects of either manipulation alone. This additive enhancement occurs without compromising the primary conditioned memory (Fig. 5H), suggesting that encoding variability and temporal spacing may engage complementary pathways underlying memory generalization. Using the protein synthesis inhibitor cycloheximide (CXM), we find that while the 24-hour conditioned memory after spaced training is strongly reduced, which confirms its reliance on classical long-term memory pathways^7,20^, the generalization enhancement triggered by encoding variability remains largely preserved (Supplementary Fig. S10).

Together, the biological findings in *Drosophila* are consistent with the computational results in ANNs. Both appropriately spaced training intervals and increased trial-to-trial sensory variability can significantly and lastingly enhance the generalization of memories. These results demonstrate a convergent computational principle of biological and artificial systems, where structured exposure to temporal or contextual variations promotes the formation of more robust and generalizable representations.

## DISCUSSION

In this study, we identify an encoding-variability-inspired principle^26–28^ as a shared computational principle that enhances generalization in both biological and artificial systems. Specifically, we demonstrate that appropriately spaced exposure to input or innate variations significantly improves generalization from training to test in ANNs (Fig. 2 and Fig. 3). This effect is robust across a variety of benchmark datasets, network architectures, and temporal ensemble strategies. Importantly, we find that the two key factors, variation strength and spaced interval, independently and additively enhance generalization. Beyond artificial systems, behavioral experiments in *Drosophila* reveal similar benefits under spaced exposure to variable training cues, showing that both spaced training intervals (Fig. 4) and increased sensory variations (Fig. 5) enhance memory generalization in olfactory associative learning. Together, these findings demonstrate how computational principles derived from artificial neural networks can guide biological experimentation, suggesting that encoding variability may operate as a unified mechanism that enhances generalization across both artificial and natural learners.

A central concept in our work is *generalization*. In machine learning, this typically refers to a model’s ability to apply knowledge from training data to unseen test data drawn from the same underlying distribution^1^. In neuroscience, although animals are often tested with the same sensory cues presented during training, it is rarely possible to exactly reproduce the internal states or environmental conditions across training and testing episodes. As a result, what is traditionally studied as *precise memory* in neuroscience implicitly involves a form of “generalization”^4,60^.

While the spacing effect has classically been linked to protein synthesis-dependent longterm memory, our results suggest that memory persistence and memory flexibility can show different sensitivities to protein synthesis inhibition. Under CXM treatment, conditioned memory after spaced training is strongly reduced, whereas the generalization enhancement induced by sensory variability remains largely preserved (Fig. S10). This pattern differs from previously reported “rapid generalization” mechanisms^61^, which rely on prior associative linking between environments. Instead, our results align with a consolidation of variable experiences that, under our experimental conditions, appears less sensitive to protein synthesis inhibition. Nonetheless, we acknowledge that a more potent or prolonged inhibition might potentially impair generalized memory as well^62^. This suggests that while temporal spacing is essential for memory stabilization, the “flexibility” of memory can be modulated by sensory variability, potentially through distinct molecular or circuit-level changes. Therefore, biological learning may utilize parallel mechanisms to balance memory specificity and generalization, a principle that aligns with the dual requirements of adaptive learning systems in both biological and computational contexts.

Notably, the enhancement of generalized memory emerges after 24 hours but not at 3 minutes post-training (Fig. 4), suggesting that spaced learning facilitates the delayed consolidation and gradual refinement of acquired knowledge into new, related contexts, without requiring preconfigured associations. We suggest that this type of exact recall in biological learning (despite identical cues) is more analogous to the concept of generalization in machine learning (i.e., from training data to similar test data), while the broader notion of generalization in biological learning (e.g., transferring across different cues or contexts) parallels the more difficult domain generalization tasks in machine learning (i.e., from training data to dissimilar test data). Our work bridges this conceptual gap by showing how spacing and variability jointly support generalization in both fields.

The spacing effect has long been recognized in neuroscience for its beneficial impact on memory formation, but its computational implications for AI remain underexplored. Our results demonstrate that incorporating spacing mechanisms, particularly through temporal ensemble strategies that mimic the temporally evolving nature of biological learning, can substantially en-hance test performance in ANNs. Unlike traditional EL methods that require training multiple models in parallel, temporal ensemble strategies efficiently aggregate internal variations over time. By further introducing structured spacing into these methods, we unlock new performance gains while maintaining computational efficiency. These insights have practical implications for a range of AI challenges, including few-shot learning^63–65^, continual learning^66–69^, and adversarial robustness^70,71^, where generalization from limited or noisy data is critical.

From the perspective of biological learning, prior studies have often manipulated either variation strength or spaced interval in isolation. For instance, encoding variability is typically studied by altering environmental conditions while keeping temporal spacing fixed, whereas spacing effects are evaluated by varying inter-trial intervals with constant stimuli^26–28^. In our ANNs experiments, training proceeds continuously, and the “spaced interval” is implemented computationally as an interval along the training steps or epochs between ensemble updates (e.g., changing cutout size, modulating dropout probability, aggregating network states, or updating teachers in KD). This design allows joint manipulation of variation strength and spaced interval to study their combined effects on generalization. Inspired by temporal ensemble strategies^30,31,33,34^, this dual modulation provides a more comprehensive understanding of the importance of encoding variability. The observed complementary effect between variation strength and spaced interval highlights their distinct, additive benefits to generalization, which may inform future designs of biological learning and memory experiments.

A limitation of our current formulation is that we primarily characterize variations through a scalar variation strength *k* with a spaced interval *s*. In practice, variations are often multidimensional, and their detailed *structure* (e.g., specific masking patterns or teacher–student pairings) may further influence diversity and generalization. A systematic analysis of how structured variations interact with the spaced interval is an important direction for future work. While our computational experiments demonstrate robust performance gains, we acknowledge that our claim linking spacing to generalization via encoding variability in ANNs remains primarily grounded in performance-level observations. Future work should incorporate more direct analyses quantifying representational diversity and internal network dynamics to further strengthen this mechanistic interpretation. In addition, while we emphasize an encoding-variability-inspired perspective for its connection to generalization and temporal ensembling, deficient-processing and study-phase retrieval mechanisms may also contribute to the spacing effect depending on task and memory phase. Furthermore, while our biological experiments demonstrate the behavioral benefits of training variability, we have not yet directly characterized the underlying coding variability at the neuronal level. Our current findings provide a bio-inspired, computational-to-behavioral validation of the encoding variability theory, but the specific neural dynamics, such as how synaptic ensembles or population activities evolve across spaced intervals, require further investigation across multiple biological levels.

Looking ahead, this study exemplifies the promise of NeuroAI^1,3,4^, an emerging field that seeks to accelerate progress in both neuroscience and AI through mutual inspiration. Our findings demonstrate how biologically grounded learning principles such as spacing and encoding variability can be translated into algorithmic gains for ANNs, while at the same time, controlled experiments in ANNs can generate testable hypotheses about biological learning mechanisms. Promising future directions include developing adaptive spacing schedules that dynamically adjust based on learning progress, extending multi-modal generalization across visual, auditory, and other sensory modalities, and integrating bio-inspired plasticity rules to further align ANNs with the dynamics of biological learning and memory. These directions may pave the way toward a unified and mechanistic understanding of generalization across both biological and artificial systems.

## METHODS

### Spacing Effect of Temporal Ensemble

We investigate how the spacing effect in biological learning can be computationally leveraged to enhance generalization in ANNs via temporal ensemble strategies. Specifically, we instantiate this principle in three representative methods, WA, dropout, and KD, which operate at the synaptic, neuronal, and network levels, respectively. Our central hypothesis is that the generalization benefits observed in biological systems through spaced repetitions and encoding variability have computational analogues in ANNs, wherein variation strength and spaced interval jointly shape learning outcomes.

#### Problem Formulation

Let 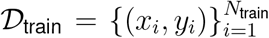 and 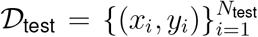 denote the training and test sets drawn from the same underlying distribution, where each sample consists of an input-label pair. A neural network model *f*_*θ*_(*x*) parameterized by *θ* is trained to minimize a supervised loss, defaulting to the cross-entropy (CE) loss for classification tasks:

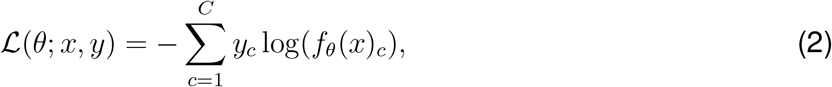

where *f*_*θ*_(*x*)_*c*_ is the predicted probability for class *c*, and *y*_*c*_ is a one-hot encoded ground-truth label.

The model’s generalization performance 𝒢 is evaluated on 𝒟_test_ using the average top-1 accuracy:

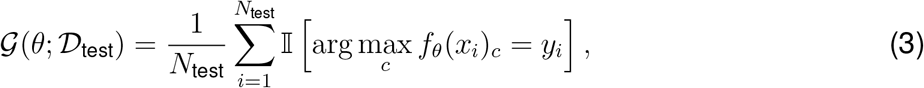

where 𝕀 [·] is the indicator function, returning 1 if the predicted class matches the true label, and 0 otherwise. *f*_*θ*_(*x*_*i*_)_*c*_ denotes the predicted probability for class *c* on test input *x*_*i*_.

#### Unified Temporal Ensemble Framework

Building upon prior theoretical efforts on temporal ensemble strategies^33,34,44^, we introduce a unified formulation that captures the core components of spacing effect with two variables: *spaced interval s* and *variation strength k*. The ensemble update process can be summarized as:

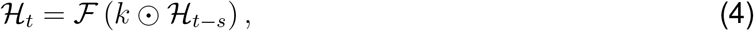

where ℋ_*t*_ denotes the internal state at the current training step *t*, which may refer to neuron activations *h*_*t*_, network parameters *θ*_*t*_, or model outputs 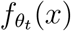 given input *x*. The operator ⊙ represents element-wise modulation. ℱis an ensembling function adapted to each strategy. In general, *k* may represent a structured operator (e.g., a mask or transformation) rather than a single scalar. Throughout the paper, we use “variation strength” as a scalar summary to enable systematic comparisons, while the detailed structure of variations may further modulate diversity and performance. The following sections define these instances and describe how the bio-inspired spacing effect is implemented in each case.

#### Dropout

Dropout^35^ operates at the neuronal level by introducing stochastic perturbations to the neuronal activations. Let *ĥ*_*t*_ denote the pre-activation vector at training step *t*. In the standard dropout formulation, a binary dropout mask *r*_*t*_ ∼ Bernoulli(*p*_*t*_) is sampled element-wise with dropout rate *p*_*t*_ ∈ [0, 1], and applied as:

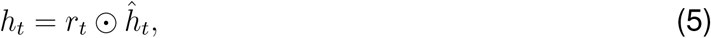

where *h*_*t*_ denotes the post-dropout neuronal activations and ⊙ denotes element-wise multiplication.

In our unified framework, the internal state at training step *t* is defined as ℋ_*t*_ = *h*_*t*_, with temporal ensembling introduced throughout training:

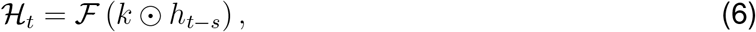

where *s* is the spaced interval, *k* corresponds to the dropout mask *r*_*t−s*_ from a previous training step, and ℱ (·) is a strategy-specific ensembling function (e.g., averaging or consistency regularization).

To further exploit the bio-inspired spacing effect, we modulate the dropout rate *p*_*t*_ periodically throughout training (Supplementary Algorithm 1). Specifically, we define:

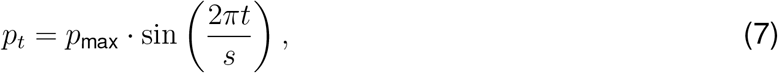

where *p*_max_ ∈ [0, 1] is the maximum dropout rate, and *s* controls the period of modulation. This introduces structured variability in the dropout-induced representations, enabling both temporal spacing and controlled variation strength to improve generalization.

#### Weight Averaging

WA^36^ operates at the synaptic level by aggregating parameter snapshots throughout training. Let 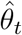 denote the original parameters updated by gradient descent at training step *t*, and *θ*_*t*_ represent the ensembled parameters. In the standard EMA formulation, the ensemble is updated iteratively as:

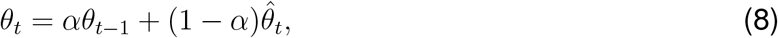

where *α* ∈ [0, 1] is the momentum coefficient controlling the influence of past parameters. The underlying base parameters evolve via standard gradient descent:

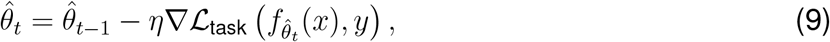

with learning rate *η* and task loss ℒ_task_.

In our unified framework, the internal state is defined as ℋ_*t*_ = *θ*_*t*_, and the ensembling process with spacing becomes:

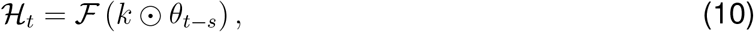

where *s* is the spacing interval, *k* corresponds to the momentum coefficient, and ℱ (·) denotes the EMA update.

To incorporate the spacing effect, we propose a spaced version of EMA (Supplementary Algorithm 2) in which parameter aggregation occurs every *s* steps:

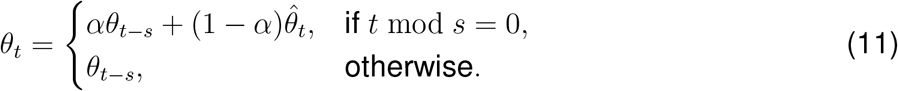

This modification introduces explicit temporal spacing into the ensemble dynamics, allowing the model to integrate more diverse parameter states and improving robustness via spaced accumulation.

#### Knowledge Distillation

KD^38^ operates at the network level by guiding a student model with soft outputs of more advanced teacher models. Among commonly-used KD methods, online KD and self KD naturally align with the concept of temporal ensembling and are suitable for investigating the bio-inspired spacing effect.

Let 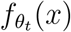 denote the teacher’s output at training step *t* for input *x*, and 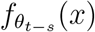 denote the student’s output from an earlier checkpoint. In standard online KD, the teacher is a more advanced model (e.g., a later snapshot or moving average), and the student attempts to match the teacher’s softened outputs. The distillation loss is defined as:

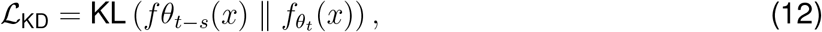

where KL(· ∥ ·) denotes the Kullback-Leibler divergence between teacher and student outputs. The student is trained using a combination of task and distillation losses:

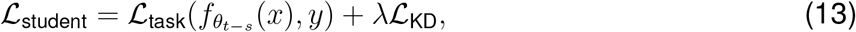

where *λ* is a weighting coefficient balancing task supervision and distillation guidance.

In our unified framework, we define the internal state at time step *t*−*s* as _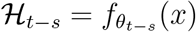_. The spaced version of online KD introduces temporal ensembling through delayed teacher-student supervision:

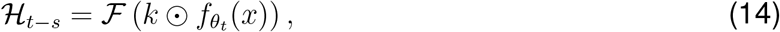

where *s* is the spaced interval, *k* corresponds to a variation operator modulating the teacher signal (e.g., temperature scaling or input perturbation), and ℱ(·) is the distillation function (e.g., KL divergence or consistency regularization).

To enforce spacing explicitly, the teacher is updated every *s* steps based on task supervision:

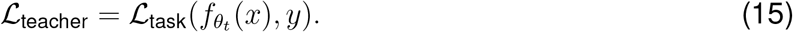

This temporal offset between teacher and student promotes representational diversity and prevents premature convergence to local minima.

For self KD^40^, the teacher and student reside within the same network, typically across different layers, heads, or temporal branches. The spaced version of self KD (Supplementary Algorithm 4) alternates which internal representations serve as distillation targets over time. The formulation remains consistent in our framework, with spacing applied to internal states rather than model snapshots, allowing earlier components to imitate more refined representations from future computation stages.

### Experimental Setup

To evaluate the impact of bio-inspired spacing effect on learning and generalization, we design parallel experiments in both biological and artificial systems.

#### Experiments with ANNs

To simulate input variability analogous to environmental changes in biological experiments, we first employ cutout augmentation^42^ on a 4-layer CNN architecture. We then extend our study by incorporating the spacing effect into representative temporal ensemble strategies across synaptic, neuronal, and network levels.

#### Datasets and Training Protocols

We evaluate on three benchmark datasets commonly used in image classification tasks, CIFAR-10^43^, CIFAR-100^43^, and Tiny-ImageNet^72^. CIFAR-10 contains 10 classes with 50k training and 10k testing images of size 32 × 32. CIFAR-100 contains 100 classes with 50k training and 10k testing images of size 32 × 32. Tiny-ImageNet contains 200 classes with 100k training and 10k testing images of size 64 × 64. Experiments are conducted on multiple backbone architectures, including ResNet-18, ResNet-50, and ResNet-101^73^. To ensure fair comparisons, we adopt identical training protocols following prior work^40,42,74^. Unless otherwise specified, all models are trained for 80 epochs using stochastic gradient descent (SGD) with momentum 0.9, batch size 128, and a constant learning rate 0.01.

##### Weight Averaging

In standard EMA^37^, the momentum model 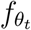 is updated at each training step from the reference model 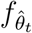. The spaced version of EMA updates 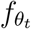 only every *s* steps, explicitly enforcing temporal spacing. The momentum coefficient *α* is set to 0.99 by default. We further implement spaced version of EMA within advanced self-supervised learning methods, including BYOL^48^, SMoG^49^ and MoCo^50^, with the official codebase https://github.com/lightly-ai/lightly. In these methods, 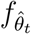 is updated using contrastive loss, and 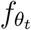 is updated using EMA.

##### Dropout

The standard dropout^35^ introduces stochasticity by randomly zeroing out neuronal activations at each training step. The spaced version of dropout instead modulates the dropout probability periodically using a sinusoidal function parameterized by the spaced interval *s* and the variation strength *k* (i.e., the maximum dropout probability *p*_max_). We set *p*_max_ = 0.3 by default. We further implement spaced version of dropout with advanced dropout variants, including DropBlock^46^, MaxDropout^45^, and Checkerboard Dropout^47^.

##### Knowledge Distillation

In standard online KD^39,75,76^, the teacher model transfers knowledge to the student model at each training step. The spaced version of online KD introduces a spaced interval *s* (in epochs), such that the teacher is trained for *s* steps before guiding the student for the next *s* steps. In the case of self KD, where deeper layers act as teachers to guide shallower ones at each training step, the spaced version alternates between training the entire network for *s* steps and then training the shallow layers for *s* steps using the deepest layers’ outputs. Following prior implementation^40,74^, we set the KD temperature to 3.0, the feature loss coefficient to 0.03, and the distillation loss coefficient to 0.3. We further implement spaced version of more advanced online KD (SHAKE^51^, CTKD^52^, LSKD^53^) and self KD (DLB^54^, PSKD^55^, TSB^56^) variants.

#### Experiments with Biological Systems

This section describes the biological experimental setup used to examine the spacing effect in *Drosophila* melanogaster.

##### Fly Strains

All fly strains are reared on standard cornmeal medium under a 12:12 hour light/dark cycle at 23^*°*^C and 60% humidity. We use wild-type flies of the *W1118* strain, which are aged 2–3 days post-eclosion and include a mixture of males and females for behavioral assays.

##### Olfactory Aversive Conditioning

To evaluate associative learning, we perform a classical Pavlovian olfactory conditioning procedure^57–59^. Flies are transferred to a behavioral room for at least 30 min to adapt to the experimental environment. Approximately 80-100 flies are subjected to the following sequential stimuli in a forward conditioning trial: air for 90 s, an odor paired with 12 pulses of 60 V electric shock (CS+) for 1 min, air for 45 s, a second odor without pairing the electric shock (CS-) for 1 min, and finally air for 45 s. For backwards training, the electric shock ends 1 min before the onset of the first CS+ odor. In the odor exposure only (OEO) control, flies are exposed to both odors in the same sequence but without any electric shock delivery. All behavioral experiments are conducted at 25^*°*^C and 60% relative humidity. The following odorants are used, all diluted in mineral oil, including 4-methylcyclohexanol (MCH, 1.0 × 10^*−*3^ dilution, Fluka), 3-octanol (3-OCT, 1.5×10^*−*3^ dilution, Sigma-Aldrich), 1-octanol (1-OCT, 2.0×10^*−*3^ dilution, J&K Scientific), and ethyl acetate (EA, 1.0 × 10^*−*3^ dilution, Alfa Aesar).

##### Drug Feeding

For pharmacological manipulation, flies are fed with 35 mM CXM (Sigma) in control solution [5% (wt/vol) glucose and 3% (vol/vol) ethanol] to block protein synthesis, while control groups (CXM-) received the vehicle solution only. To inhibit de novo protein synthesis, flies were fed 35 mM cycloheximide (CXM; Sigma-Aldrich) in vehicle solution [5% (wt/vol) glucose and 3% (vol/vol) ethanol] for 14-16 hours prior to massed or spaced training. This pharmacological treatment was maintained throughout the 24-hour retention interval until testing. This concentration and feeding protocol has been extensively validated to effectively block protein synthesis in *Drosophila* memory assays^58,77^.

##### Memory Testing and Performance Index

Memory performance is evaluated using a standard T-maze apparatus at 3 minutes (immediate memory) or 24 hours (long-term memory) after training. Flies are given 1 minute to choose between the CS+ odor (or a structurally similar analog) and a novel odor. The preliminary performance index (PI) is calculated as:

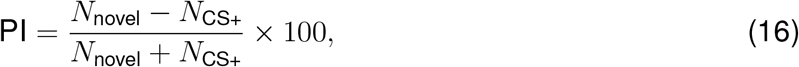

where *N*_novel_ and *N*_CS+_ denote the number of flies in the novel odor and CS+ arms, respectively. A PI of 100 indicates that all flies make the right choice to avoid the odor paired with the electric shock, while a PI of 0 indicates no memory retention, as reflected by a 50:50 distribution between the arms. To balance naive odor bias, two reciprocal groups are trained and tested simultaneously. One group is trained with CS+/CS-, and the other with odor exposure only or backwards training (i.e., shock delivered 1 minute before CS+ odor exposure). The final learning performance is calculated as a corrected PI, defined by subtracting the PI of the control group from that of the associative training group to account for non-associative components.

## Supporting information

supplemental files of our implementation

## RESOURCE AVAILABILITY

### Lead contact

Requests for further information and resources should be directed to and will be fulfilled by the lead contact, Liyuan Wang.

### Materials availability

This study did not generate new materials.

### Data and code availability

#### Data Availability

All benchmark datasets used in this paper are publicly available, including CIFAR-10/100^43^ (https://www.cs.toronto.edu/~kriz/cifar.html) and Tiny-ImageNet^72^ (https://www.image-net.org/download.php). Statistical analysis is performed using GraphPad Prism software. Data are considered normally distributed if they passed the Shapiro-Wilk test (for n*<*8) or the Anderson-Darling test (for n≥ 8). For normally distributed data, comparisons between the two groups are performed using the two-tailed unpaired t-test; comparisons between multiple groups are performed using the One-way ANOVA test followed by Dunnett’s multiple comparisons test, and the two-way ANOVA test followed by Sidak’s multiple comparisons test. Results are reported as n.s. (not significant) for p *>* 0.05, * for p *<* 0.05, ** for p *<* 0.01, *** for p *<* 0.001, and **** for p *<* 0.0001.

#### Code Availability

Our source code is available at GitHub (https://github.com/SunGL001/spacing_generalization) and has been archived at Zenodo^78^.

## ACKNOWLEDGMENTS

This work is supported by the STI2030-Major Projects (No. 2022ZD0204900 to Y.Z.), the Beijing Major Science and Technology Project (No. Z251100008425003 to L.W. and Y.Z.), the NSFC Projects (Nos. 62406160 to L.W., 32021002 to Y.Z.), and the National Science and Technology Major Project (No. 2022ZD01163013 to B.L.).

## AUTHOR CONTRIBUTIONS

Study conception and design: G.S., L.W. Computational experiment performing: G.S., H.Y. Biological experiment performing: N.H. Visualization and data analysis: G.S., N.H., H.Y., L.W. Funding acquisition: Y.Z., L.W., B.L. Results were discussed and interpreted by: G.S., N.H., H.Y., L.W., J.Z., Q.L., B.L., Y.Z. Manuscript was written by: G.S., N.H., L.W. Manuscript was discussed by: G.S., N.H., H.Y., J.Z., Q.L., B.L., Y.Z., L.W.

## DECLARATION OF GENERATIVE AI AND AI-ASSISTED TECHNOLOGIES

Large language models were used to polish the manuscript. The authors have thoroughly reviewed and edited all content and take full responsibility for the published work.

## SUPPLEMENTAL INFORMATION INDEX

**Figure S1:**
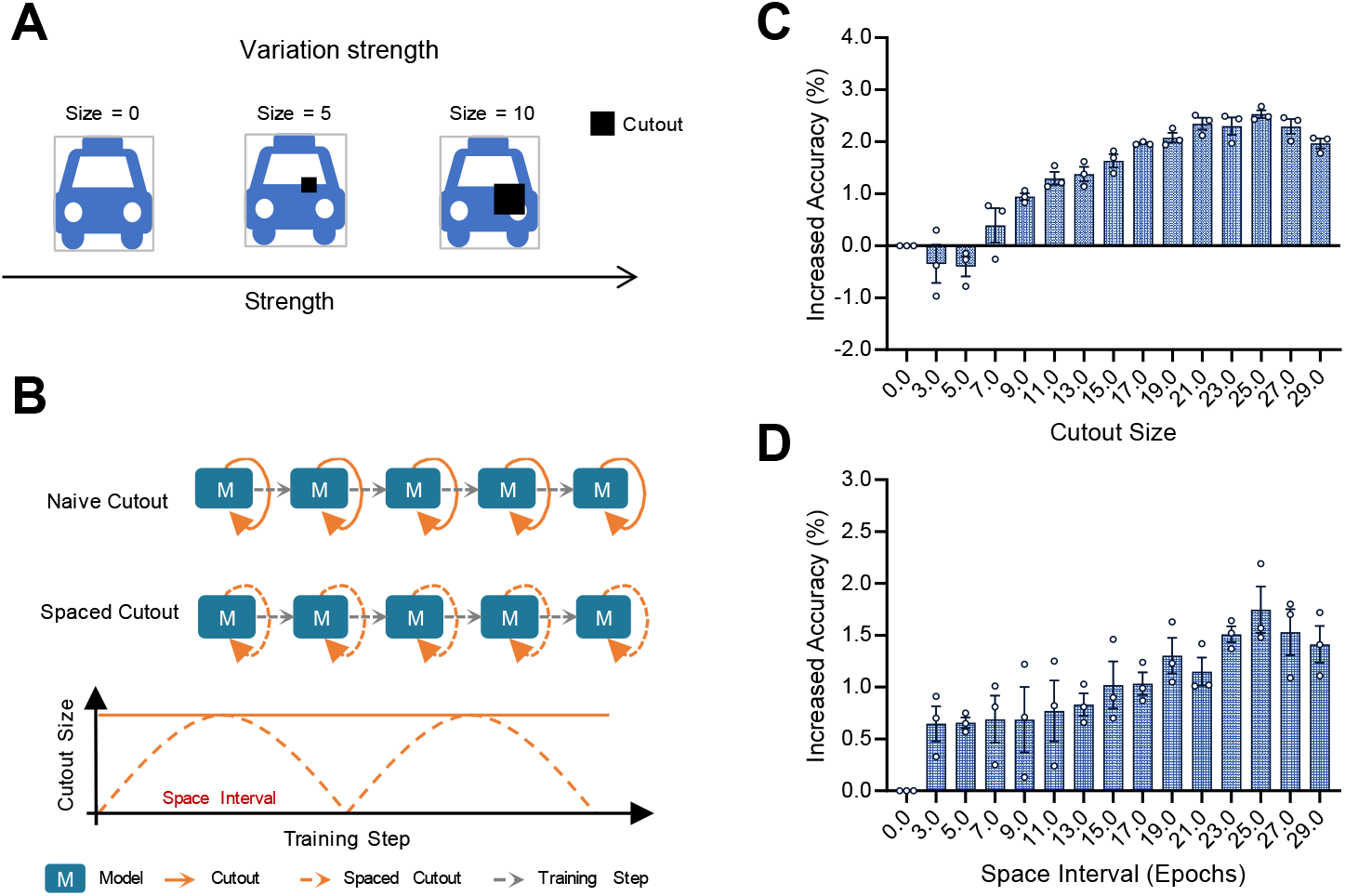
Impact of variation strength and space interval in cutout augmentation. (**A**) Variation strength is modulated by the cutout size, which controls the degree of occlusion applied to the input image. (**B**) Spaced interval determines how frequently the cutout size is altered across training epochs. (**C**) Performance gains of ResNet-18 on CIFAR-10 exhibit an inverted U-shaped trend with respect to cutout size. (**D**) Periodically varying cutout size with different spaced intervals also produces an inverted U-shaped curve. All results are averaged over three runs with different random seeds. Data are presented as mean ± SEM.

**Figure S2:**
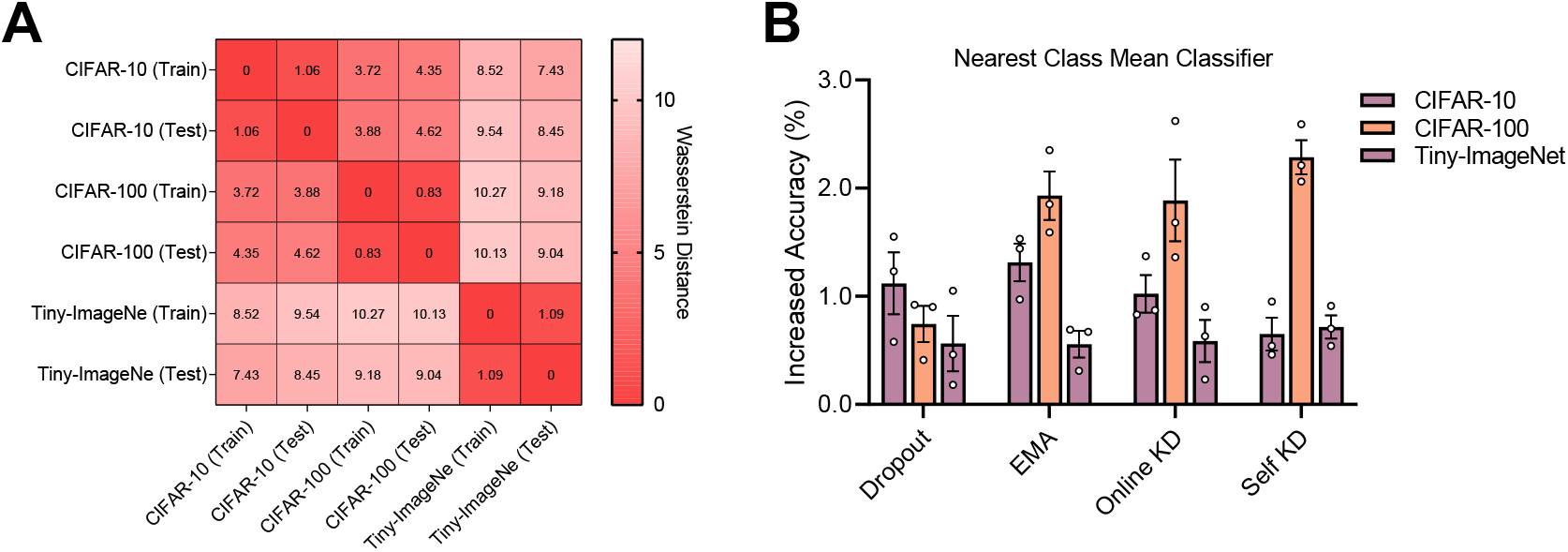
Quantitative assessment of training-test similarity and generalization. (**A**) Average data differences under the Wasserstein distance metric. The heatmap displays pairwise distances between 5,000 randomly selected samples from the training and test splits of each dataset. (**B**) Generalization of cross-dataset representations on ResNet-18. The learned representations from the training set of CIFAR-100 are evaluated with a nearest class mean (NCM) classifier on the test sets of CIFAR-10, CIFAR-100, and Tiny-ImageNet. We report the increased accuracy (%) achieved by the Spaced strategies relative to the Naive baselines.

**Figure S3:**
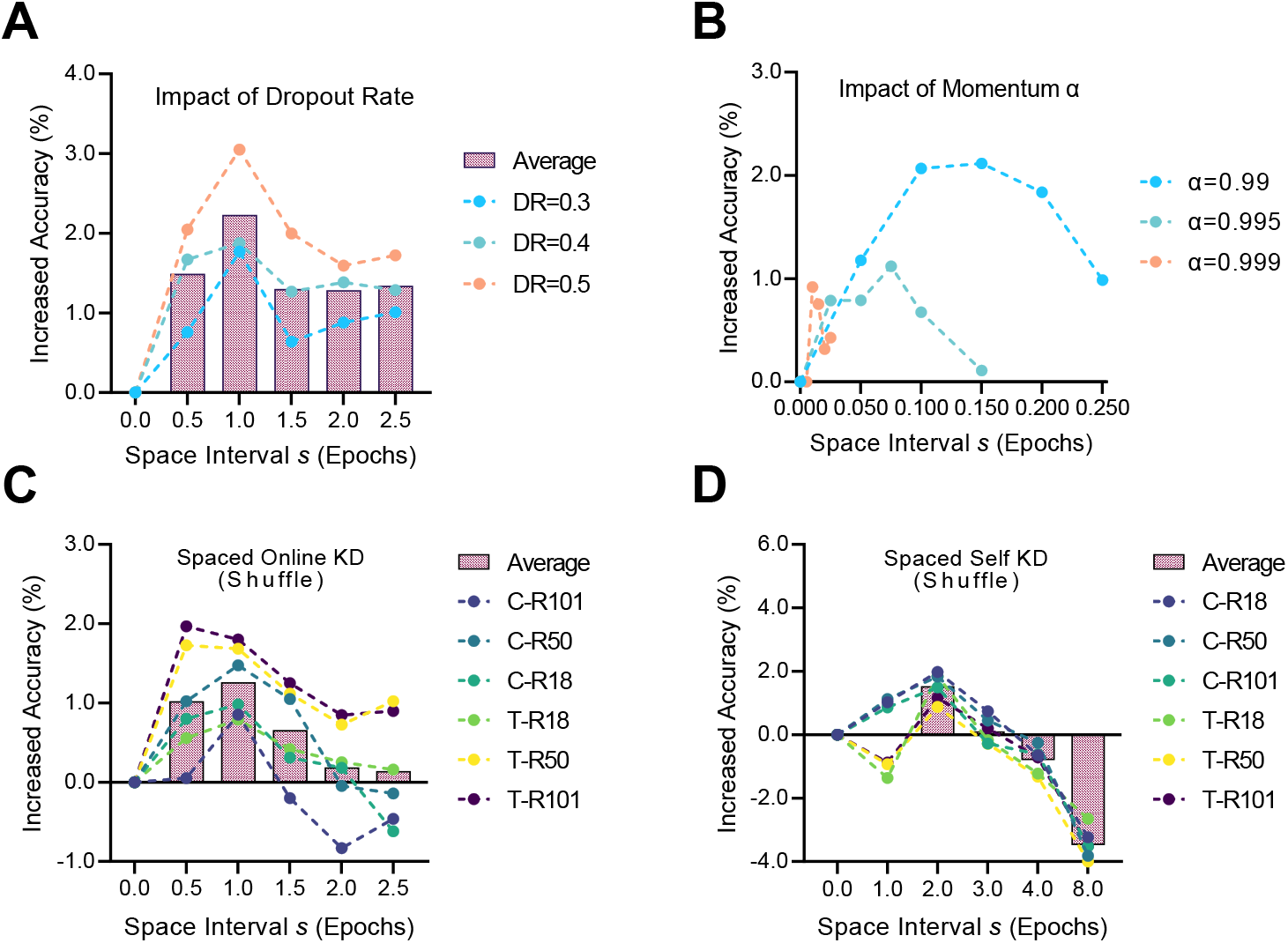
Impact of variation strength and space interval on temporal ensemble strategies. (**A**) Performance gains under different dropout rates (*p*_max_ ∈ {0.3, 0.4, 0.5}) and spacing intervals using the spaced version of dropout. (**B**) Performance gains under different momentum coefficients (*α* ∈ {0.99, 0.995, 0.999}) and spacing intervals using the spaced version of EMA. (**C**) Performance gains when teacher and student models receive shuffled training data at spaced intervals using spaced version of online KD. (**D**) Performance gains when teacher and student models receive shuffled training data at spaced intervals using spaced version of self KD. All results are averaged over three runs with different random seeds. Data are presented as mean ± SEM. See Supplementary Table S1 for the original results of baselines and spaced variants.

**Figure S4:**
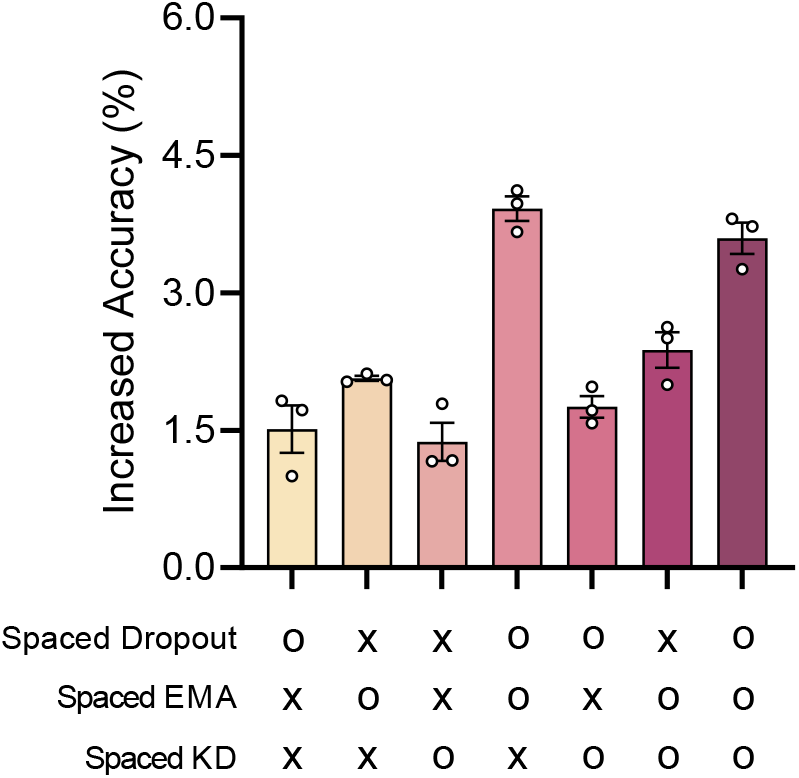
Additional benefits of combining spaced temporal ensemble strategies. We evaluate the test accuracy when combining spaced versions of dropout, EMA, and KD with ResNet-18 on CIFAR-100. “o“ indicates the strategy is enabled, while “×“ indicates it is disabled. To mitigate potential selection bias, the specific spacing intervals applied for each experiment are explicitly reported in the relevant figures and Supplementary Table S1. Data are presented as mean ± SEM over three independent runs.

**Figure S5:**
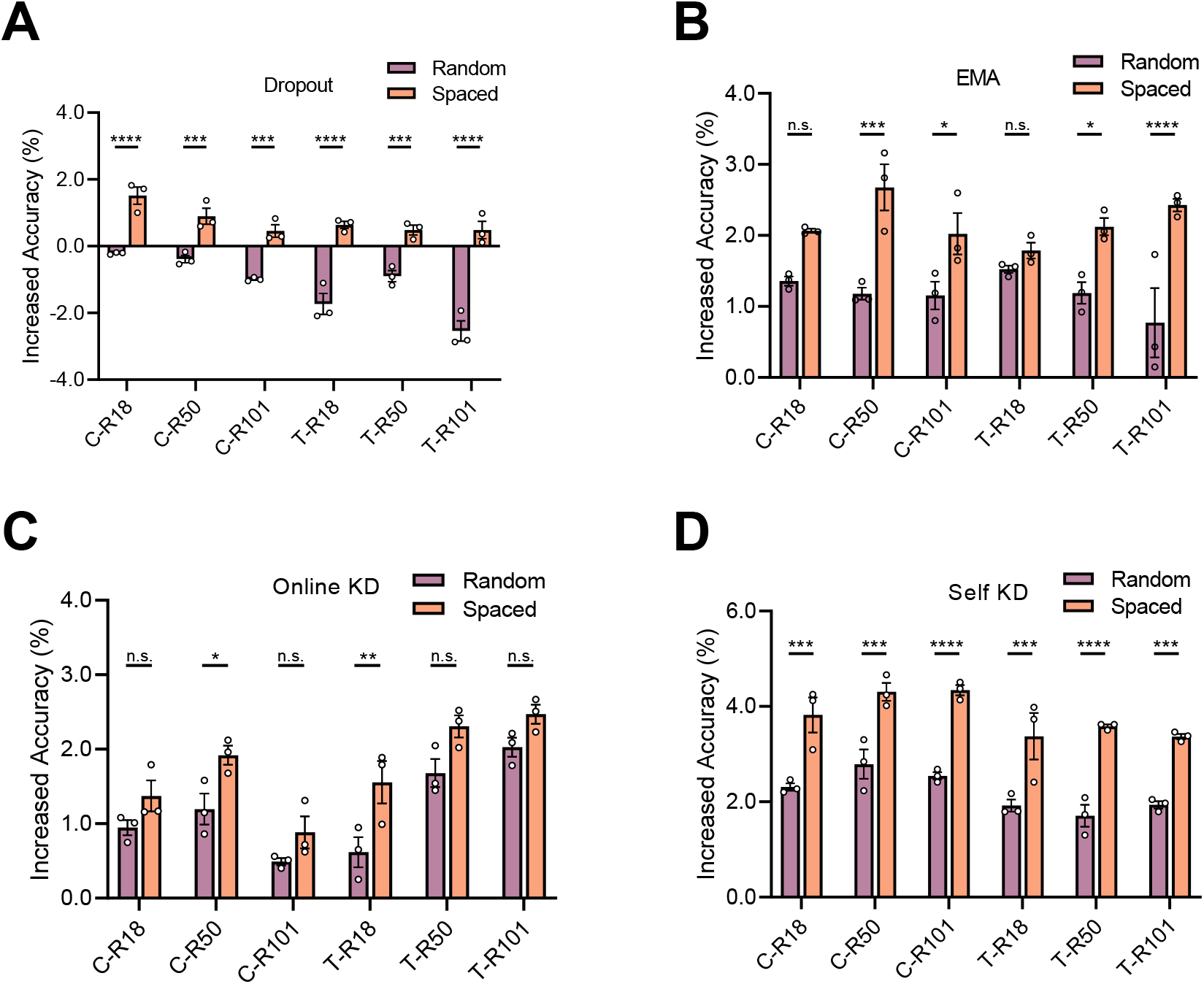
Periodicity in spacing schedules is critical for enhanced generalization. We compare the standard “Spaced” strategies (periodic intervals) against “Random” variants where the variation schedule was shuffled (irregular intervals) while maintaining the same average variation frequency. (**A**) Performance comparison for dropout. (**B**) Performance comparison for EMA. (**C**) Performance comparison for online KD. (**D**) Performance comparison for self KD. C, CIFAR-100. T, Tiny-ImageNet. R18, ResNet-18. R50, ResNet-50. R101, ResNet-101. All results are averaged over three runs with different random seeds. Data are presented as mean ± SEM. Statistical significance is determined by one-way ANOVA with Dunnett’s multiple comparisons test. *p≤0.05, **p≤0.01, ****p≤0.0001, n.s., not significant.

**Figure S6:**
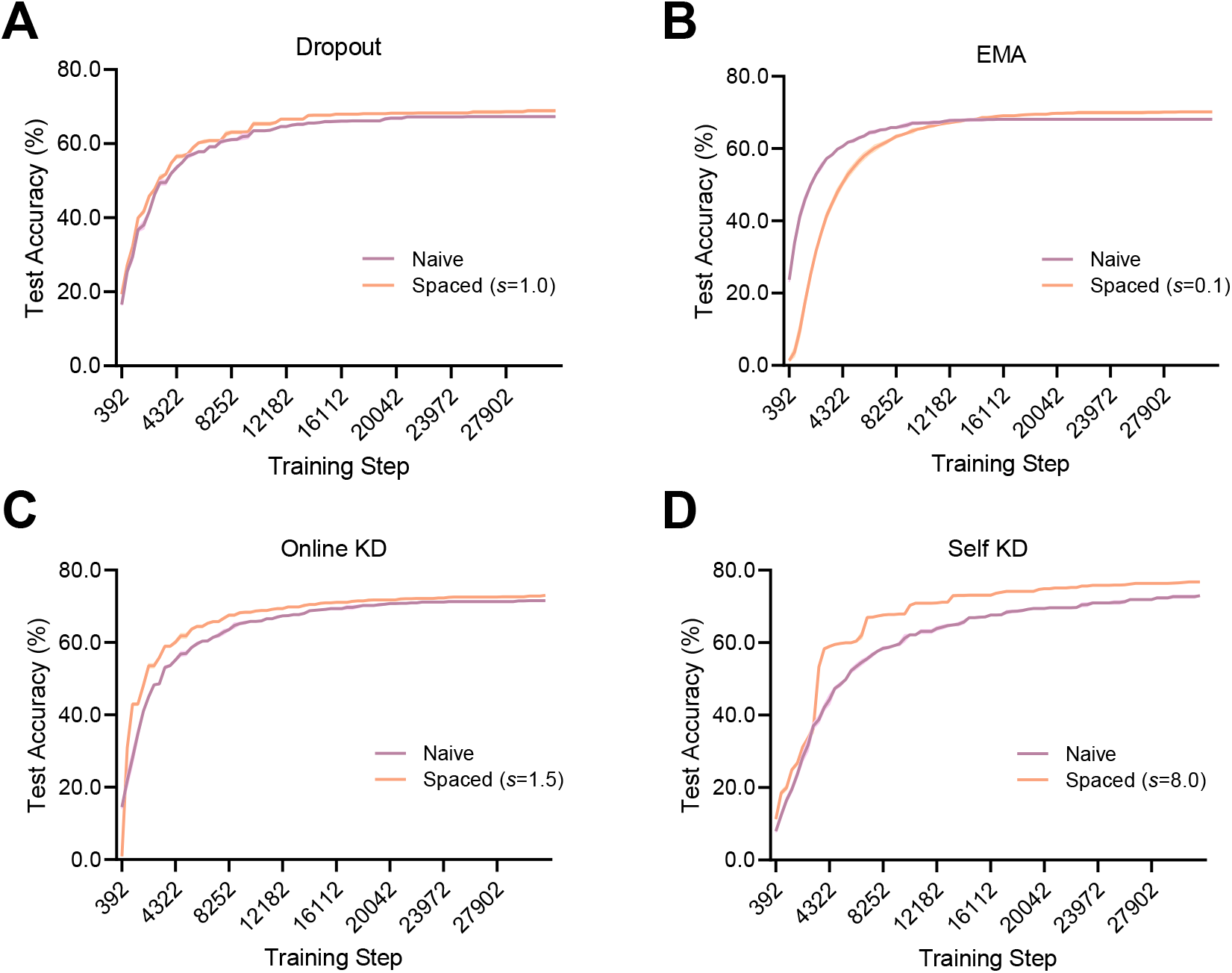
Impact of spaced training on learning dynamics and performance ceiling. We compare the test accuracy trajectories of standard “Naive” baselines versus their “Spaced” counterparts over training steps. (**A**) Training curve for dropout. (**B**) Training curve for EMA. (**C**) Training curve for online KD. (**D**) Training curve for self KD. All experiments use ResNet-18 on CIFAR-100. Data are averaged over three independent runs.

**Figure S7:**
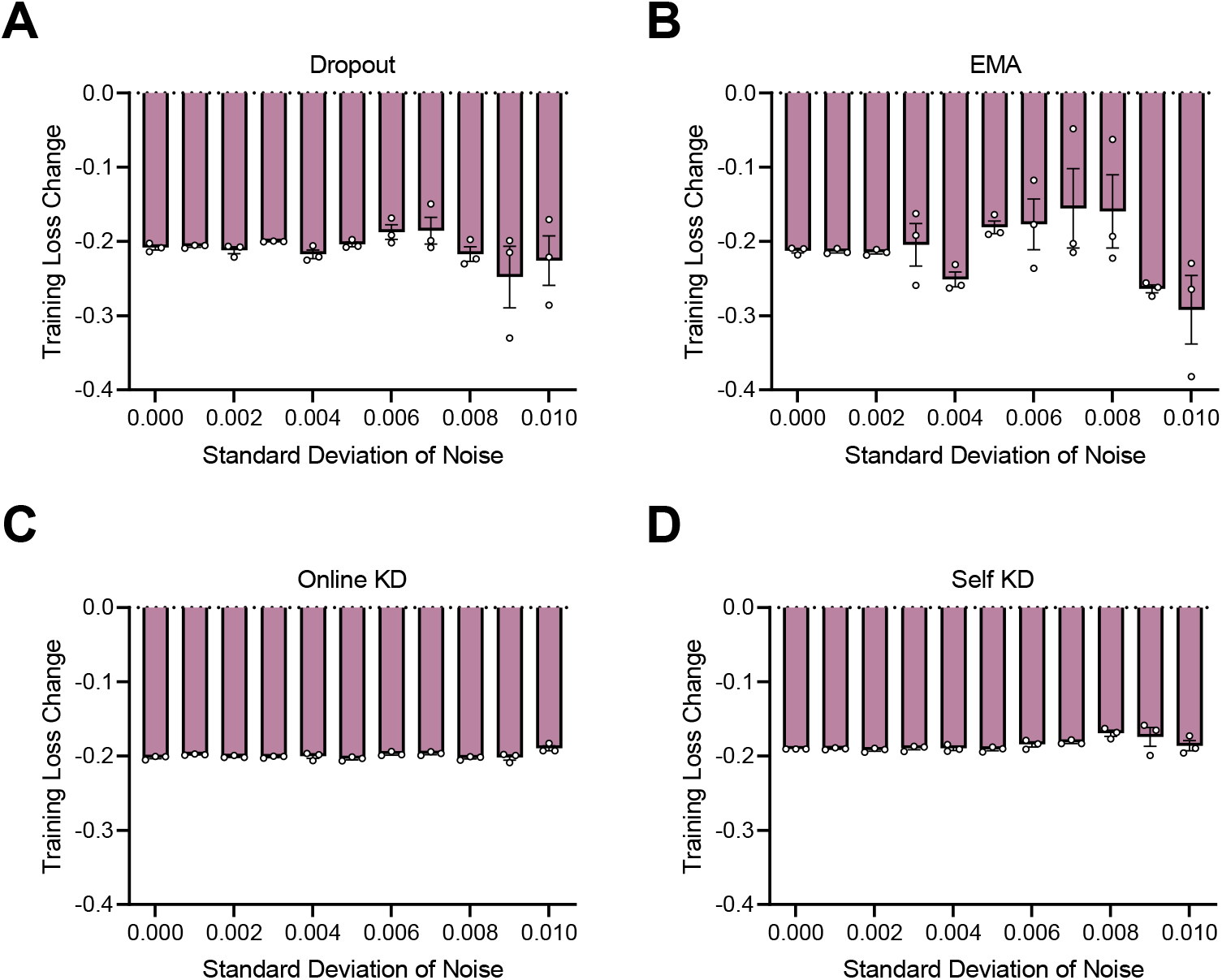
Impact of parameter perturbations with Gaussian noise. The bar display the training loss differences (Δℒ = ℒ_Spaced_ − ℒ_Naive_) between the standard “Naive” baselines and our “Spaced” counterparts. A negative value indicates that the Spaced variant experiences a smaller increase in training loss compared to the Naive baseline, suggesting a more robust parameterized solution. (**A**) Training loss differences for dropout. (**B**) Training loss differences for EMA. (**C**) Training loss differences for online KD. (**D**) Training loss differences for self KD. All experiments use ResNet-18 on CIFAR-100. Data are averaged over three independent runs.

**Figure S8:**
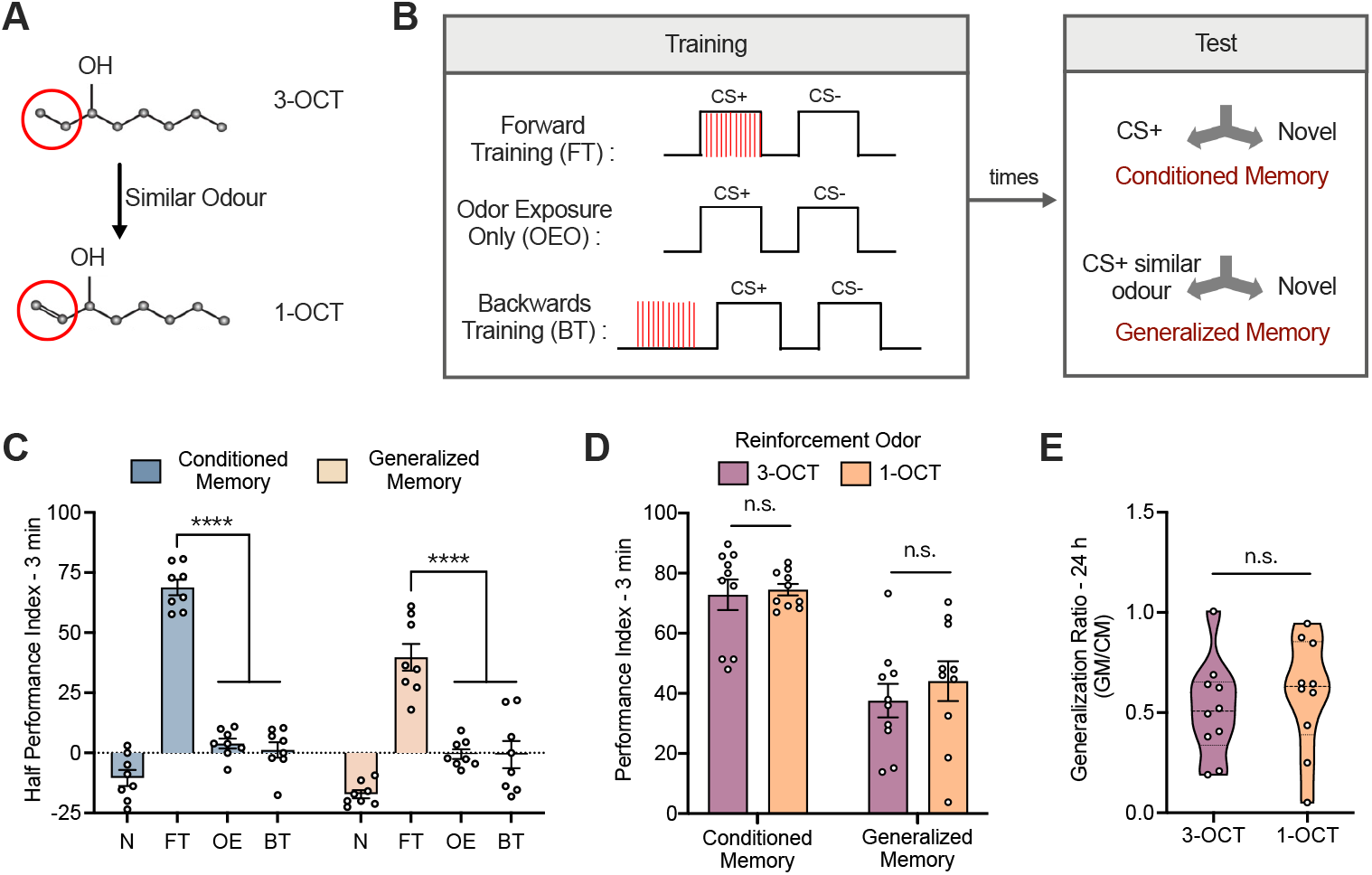
*Drosophila* exhibit generalization to a similar odor after training. (**A**) 3-OCT and 1-OCT are chemically similar odors. (**B**) Experimental paradigm for generalization training. Different groups of flies underwent forward training (FT), odor exposure only (OEO), or backwards training (BT). Conditioned and generalized memory are tested at indicated time points by presenting the CS+ odor (3-OCT) or a structurally similar odor (1-OCT), respectively, against a novel odor (MCH). (**C**) Both conditioned and generalized memory in the FT group are significantly higher than in control groups at 3 min. n=8. (**D**) Altering the reinforcement odor does not affect conditioned or generalized memory at 3 min. n=10. (**E**) Altering the reinforcement odor does not affect the generalization ratio at 3 min. n=10. Data are presented as mean ± SEM. Statistical significance is determined by two-way ANOVA with Sidak’s multiple comparisons test or unpaired t test. ****p≤0.0001, n.s., not significant.

**Figure S9:**
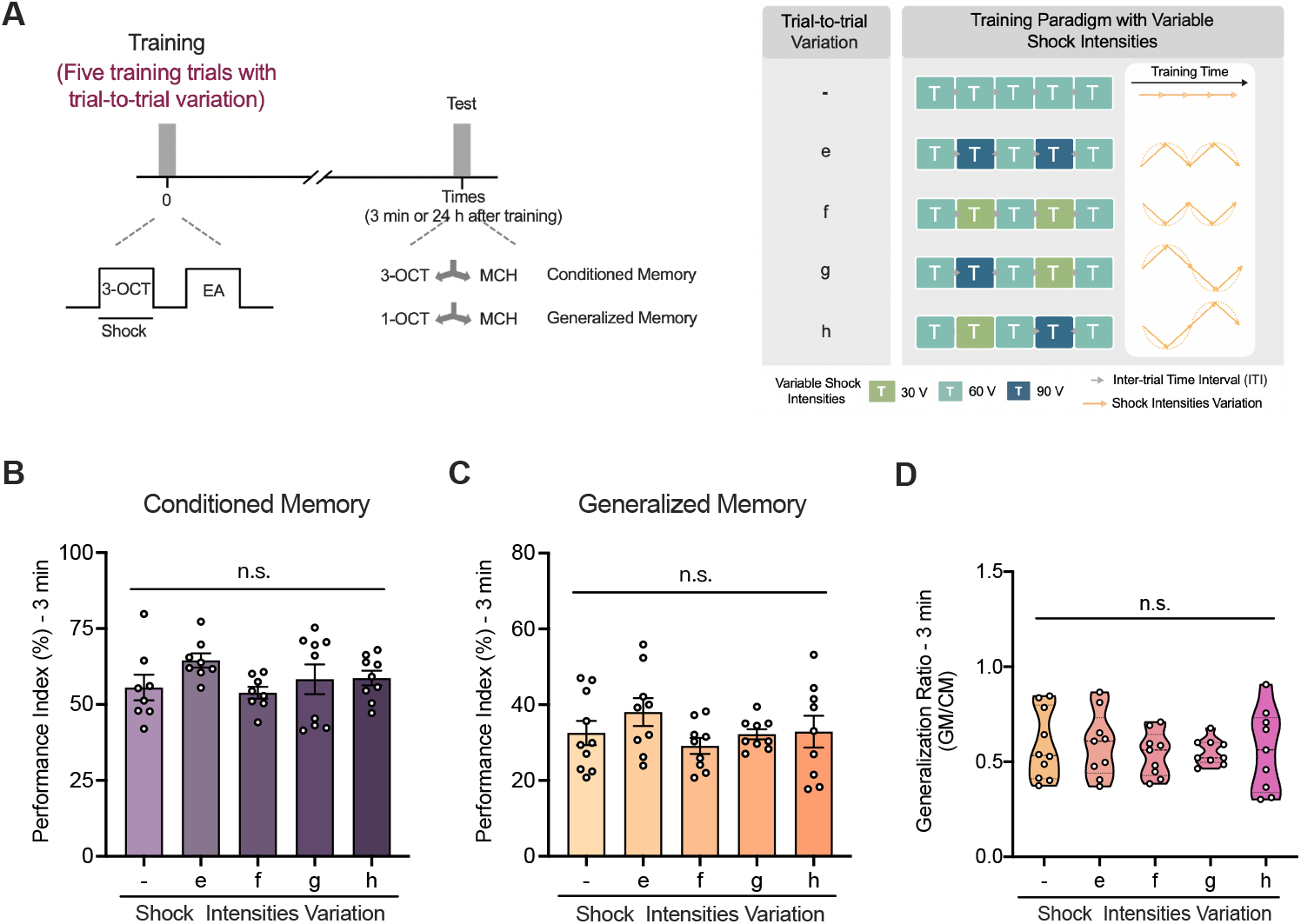
Massed training with variable shock intensity has no effect on memory. (**A**) Experimental paradigm for massed training with variable shock input. The control group (-) receives five trials with constant shock intensity, while experimental groups (e-h) receive trials with varying intensities to introduce encoding variability. (**B**-**D**) Conditioned memory (**B**, n=8-9), generalized memory (**C**, n=9-10), and generalization ratio (**D**, n=9-10) at 3 min are all no significant difference between control and experimental groups (e-h). Data are presented as mean ± SEM. Statistical significance is determined by one-way ANOVA with Dunnett’s multiple comparisons test. n.s., not significant.

**Figure S10:**
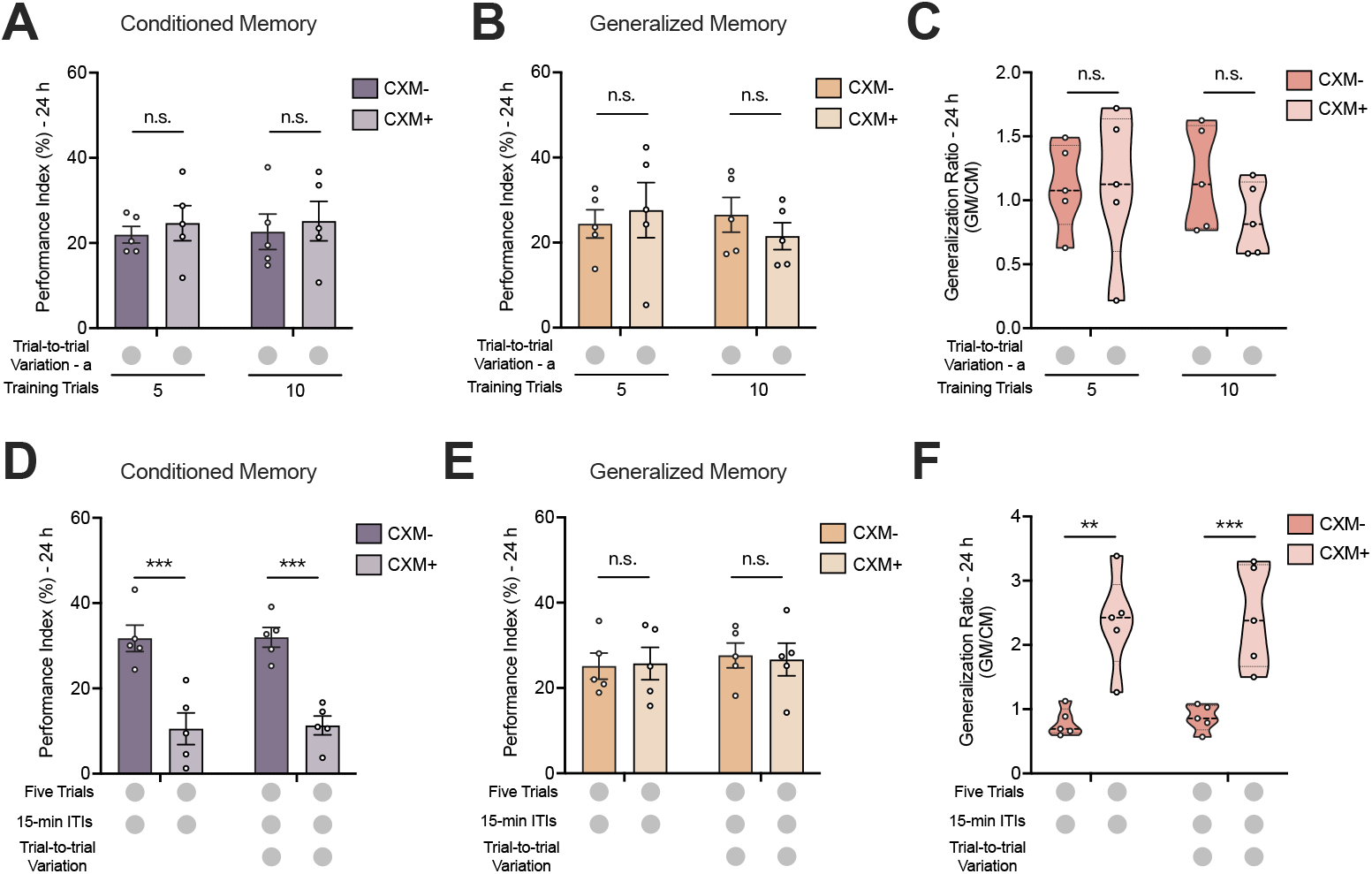
Impact of protein synthesis inhibition on memory persistence and generalization. (**A–C**) Trial-to-trial variation enhances generalization independently of protein synthesis during massed training. Conditioned memory (**A**, n=5), generalized memory (**B**, n=5), and generalization ratio (**C**, n=5) measured 24 h after 5 or 10 massed trials with trial-to-trial variation. No significant differences were observed between Control and CXM-treated groups. (**D–F**) Mechanistic dissociation between conditioned memory and generalization enhancement in spaced training. (**D**) CXM treatment significantly abolishes 24 h conditioned memory following spaced training compared to controls. n=5. (**E**) Generalized memory at 24 h remains intact in the CXM-treated group, showing no significant difference from controls. n=5. (**F**) The generalization ratio is significantly elevated in CXM-treated groups. n=5.

**Table S1:**
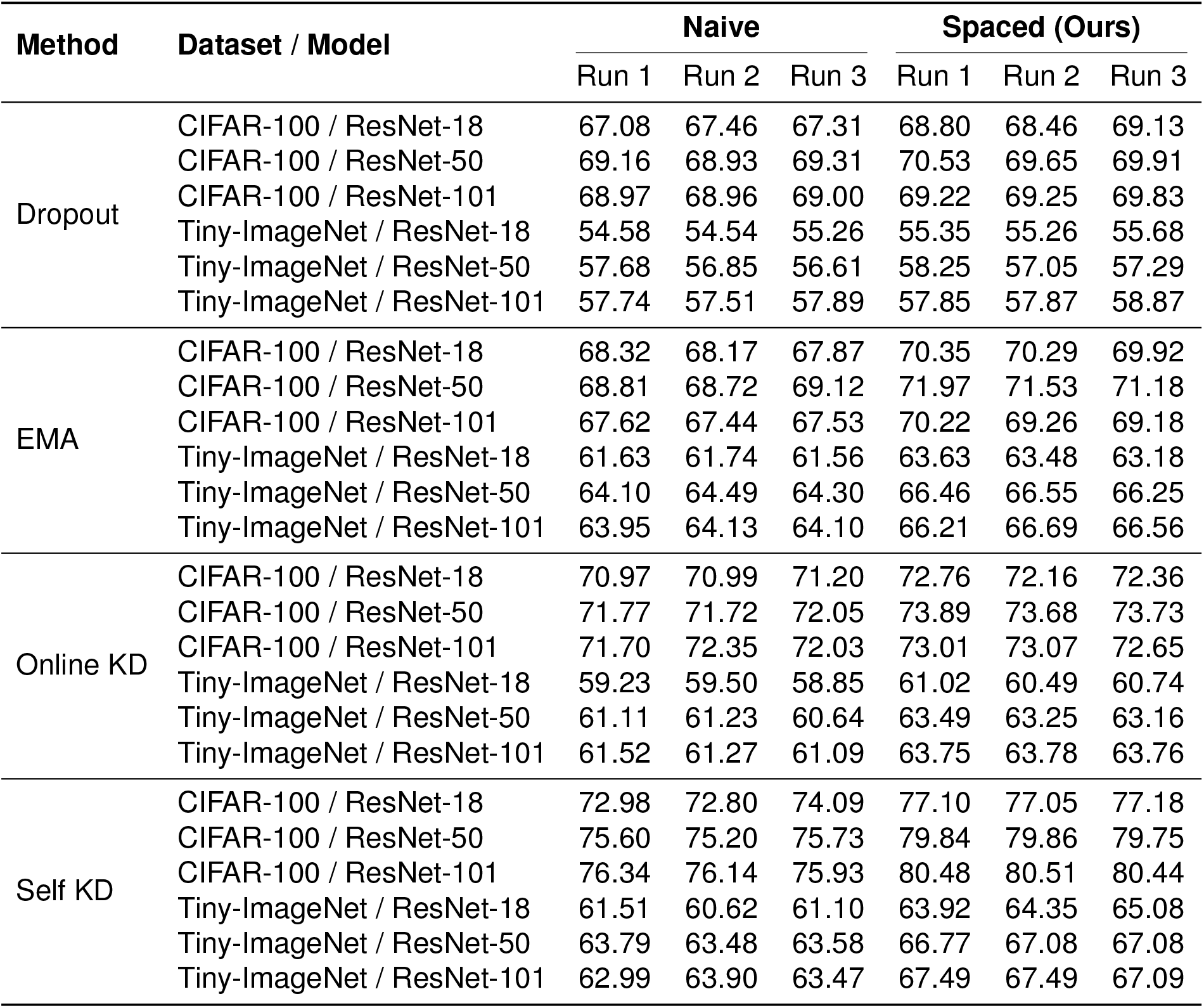
Detailed experimental results across multiple runs. This table reports the absolute test accuracy (%) for three independent runs with different random seeds. We compare the “Naive” baselines against our proposed “Spaced” variants across four temporal ensemble strategies: Dropout (Fig. 2C, *s* = 1.0), EMA (Fig. 2G, *s* = 0.1), Online KD (Fig. 3C, *s* = 1.5), and Self KD (Fig. 3G, *s* = 12.0). Experiments were conducted on CIFAR-100 and Tiny-ImageNet datasets using ResNet-18, ResNet-50, and ResNet-101 backbones. The “Spaced” variants consistently achieve higher test accuracy compared to the naive baselines across all trials and network architectures.

### Algorithm 1

Spaced version of dropout

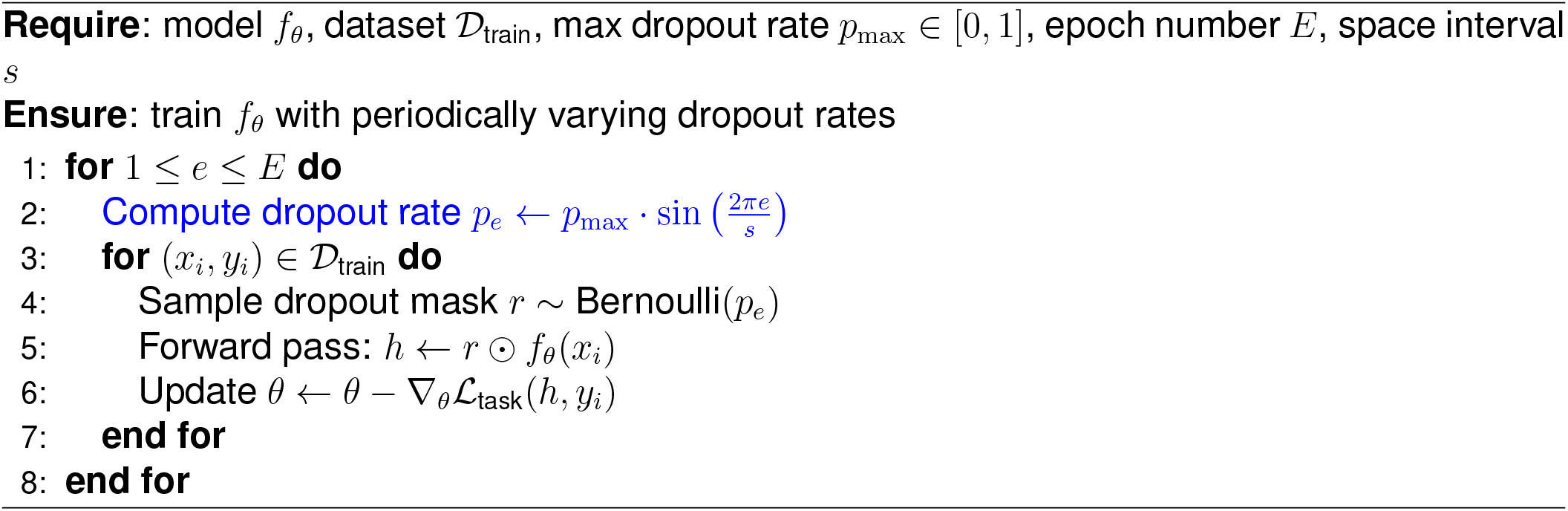

### Algorithm 2

Spaced version of EMA

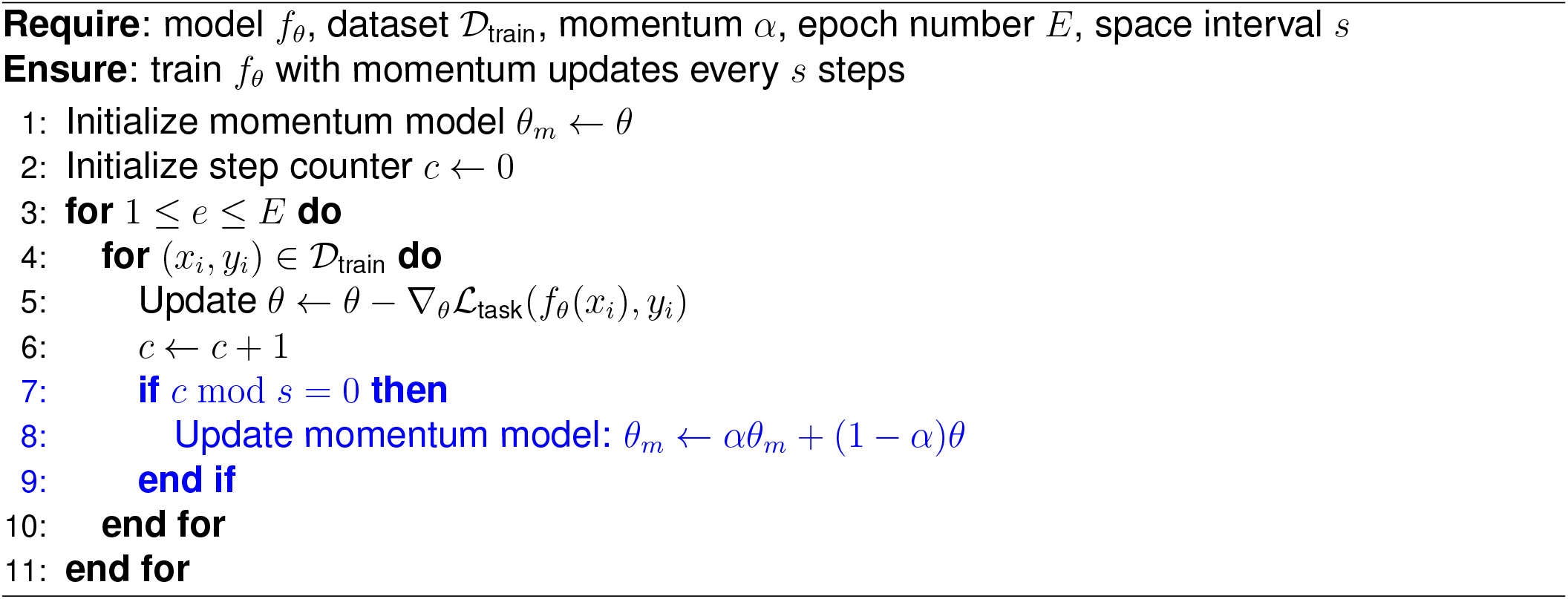

### Algorithm 3

Spaced version of online KD

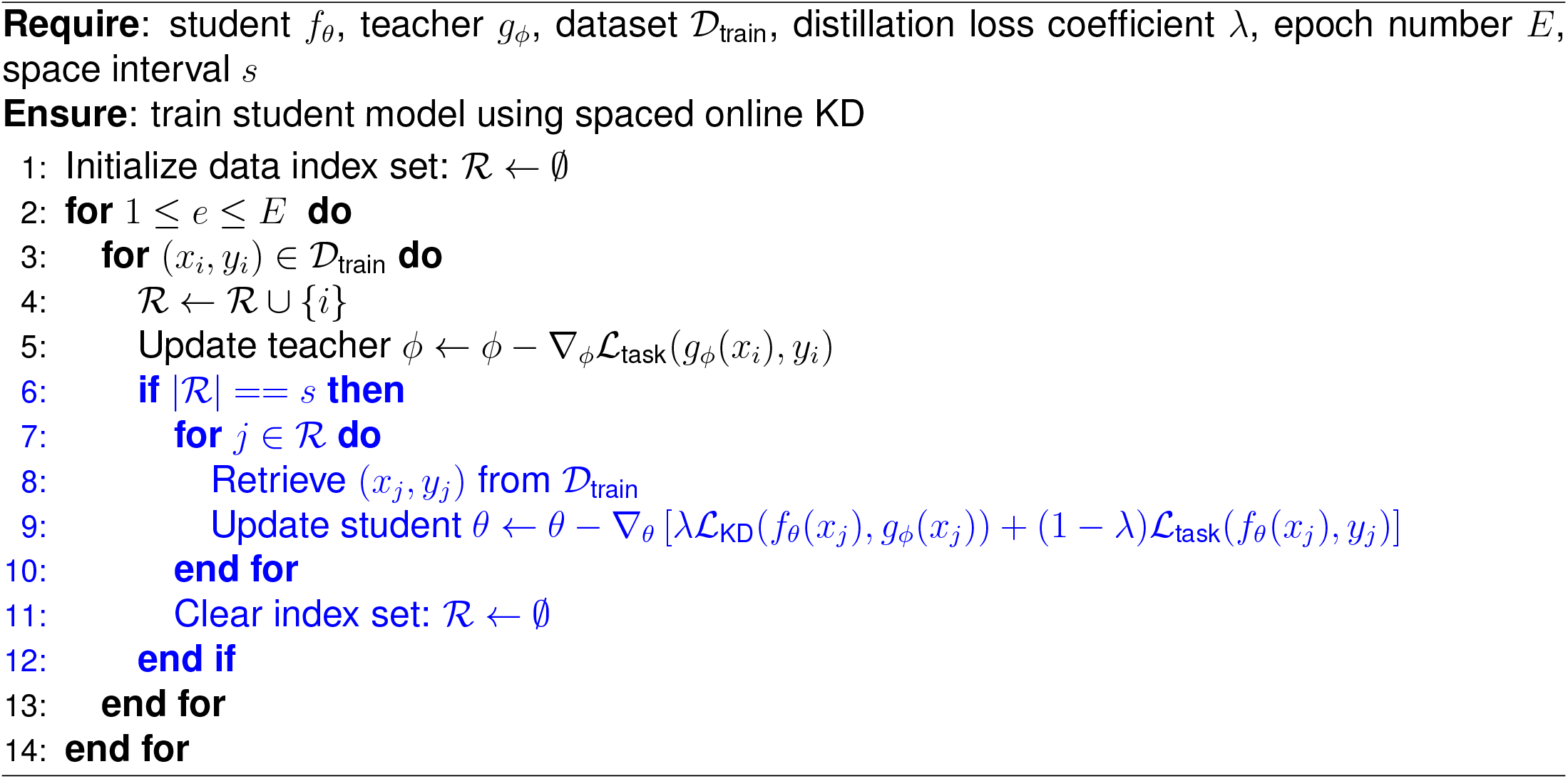

### Algorithm 4

Spaced version of self KD

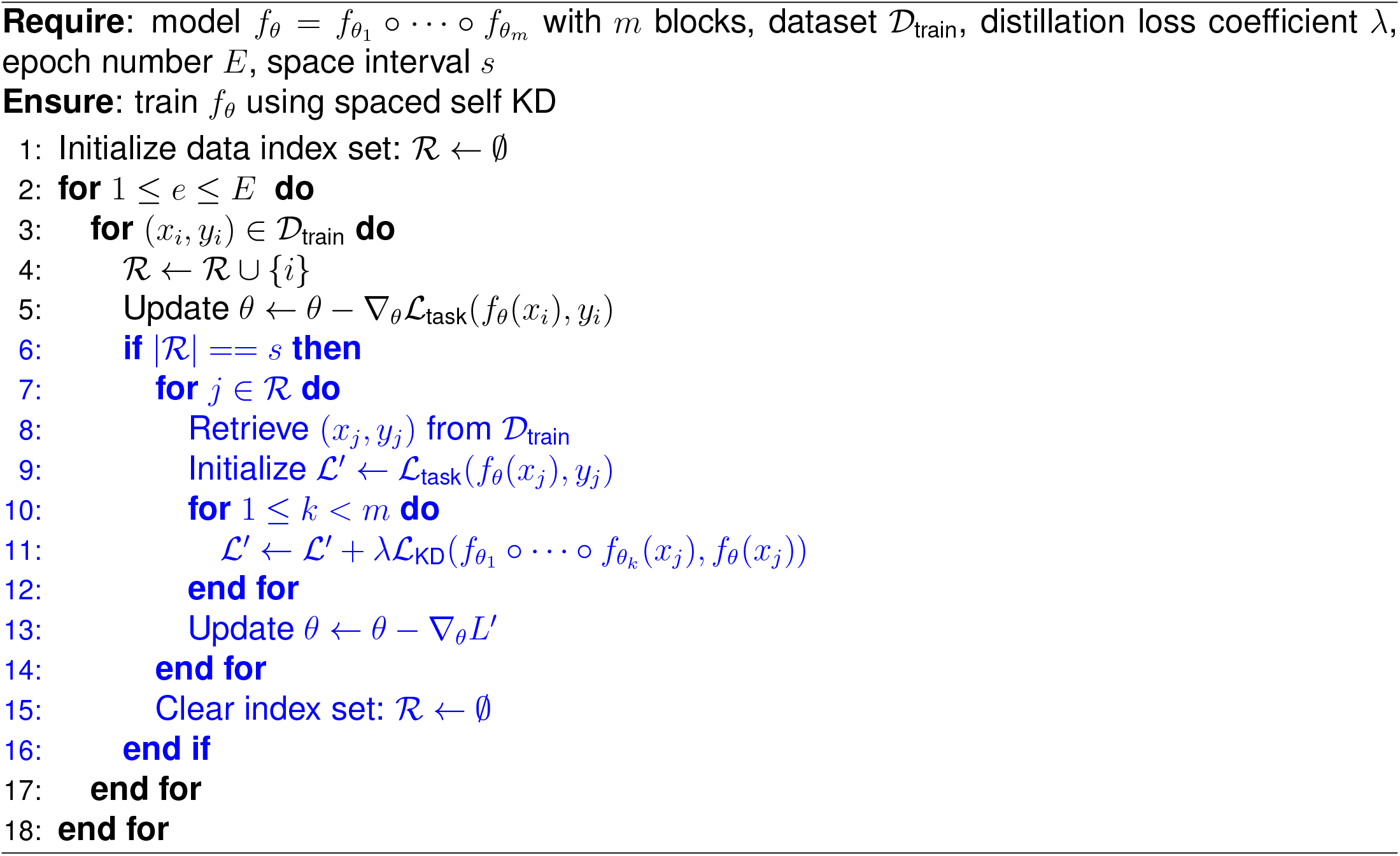

