## supplemental files of our implementation for "Spacing effect improves generalization in biological and artificial systems"

### SUPPLEMENTAL INFORMATION INDEX

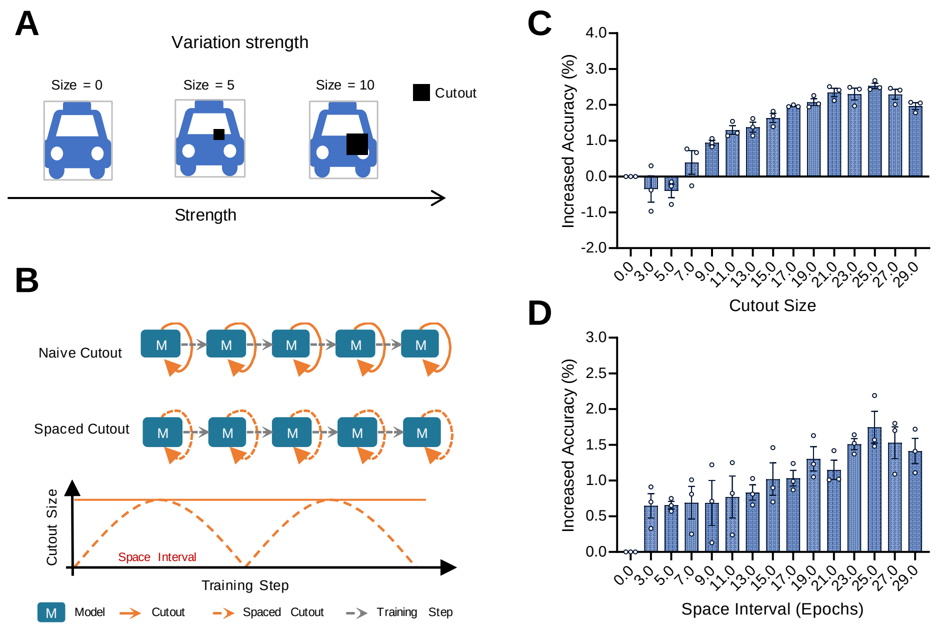


Figure S1: **Impact of variation strength and space interval in cutout augmentation.** (**A**) Variation strength is modulated by the cutout size, which controls the degree of occlusion applied to the input image. (**B**) Spaced interval determines how frequently the cutout size is altered across training epochs. (**C**) Performance gains of ResNet-18 on CIFAR-10 exhibit an inverted U-shaped trend with respect to cutout size. (**D**) Periodically varying cutout size with different spaced intervals also produces an inverted U-shaped curve. All results are averaged over three runs with different random seeds. Data are presented as mean ± SEM.

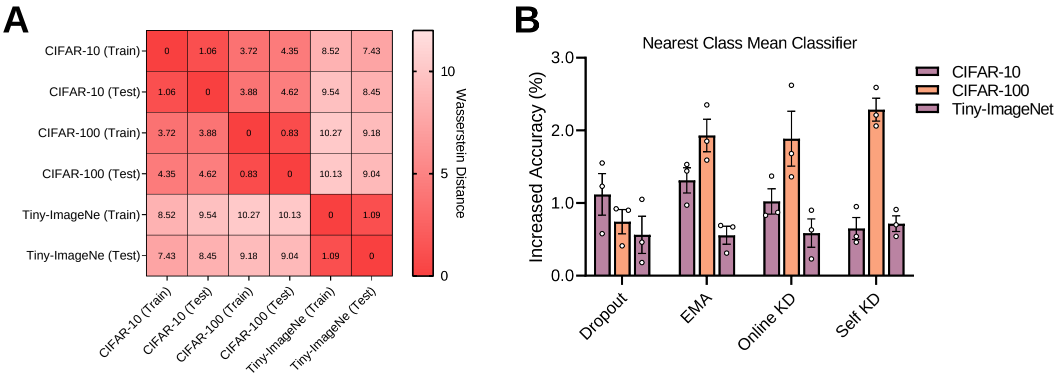


Figure S2: **Quantitative assessment of training-test similarity and generalization.** (**A**) Average data differences under the Wasserstein distance metric. The heatmap displays pairwise distances between 5,000 randomly selected samples from the training and test splits of each dataset. (**B**) Generalization of cross-dataset representations on ResNet-18. The learned representations from the training set of CIFAR-100 are evaluated with a nearest class mean (NCM) classifier on the test sets of CIFAR-10, CIFAR-100, and Tiny-ImageNet. We report the increased accuracy (%) achieved by the Spaced strategies relative to the Naive baselines.

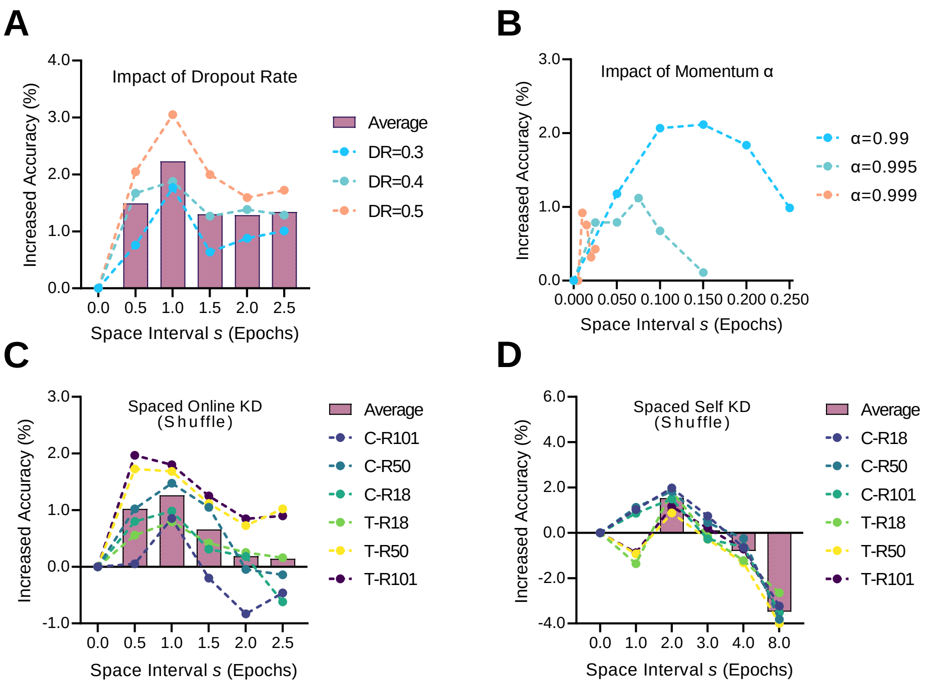


Figure S3: **Impact of variation strength and space interval on temporal ensemble strategies.** (**A**) Performance gains under different dropout rates ($p_{\max} \in \{0.3, 0.4, 0.5\}$) and spacing intervals using the spaced version of dropout. (**B**) Performance gains under different momentum coefficients ($\alpha \in \{0.99, 0.995, 0.999\}$) and spacing intervals using the spaced version of EMA. (**C**) Performance gains when teacher and student models receive shuffled training data at spaced intervals using spaced version of online KD. (**D**) Performance gains when teacher and student models receive shuffled training data at spaced intervals using spaced version of self KD. All results are averaged over three runs with different random seeds. Data are presented as mean ± SEM. See Supplementary Table S1 for the original results of baselines and spaced variants.

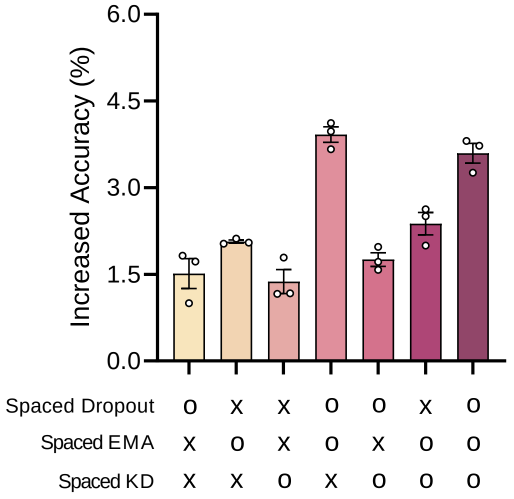


Figure S4: **Additional benefits of combining spaced temporal ensemble strategies.** We evaluate the test accuracy when combining spaced versions of dropout, EMA, and KD with ResNet-18 on CIFAR-100. "○" indicates the strategy is enabled, while "×" indicates it is disabled. To mitigate potential selection bias, the specific spacing intervals applied for each experiment are explicitly reported in the relevant figures and Supplementary Table S1. Data are presented as mean $\pm$ SEM over three independent runs.

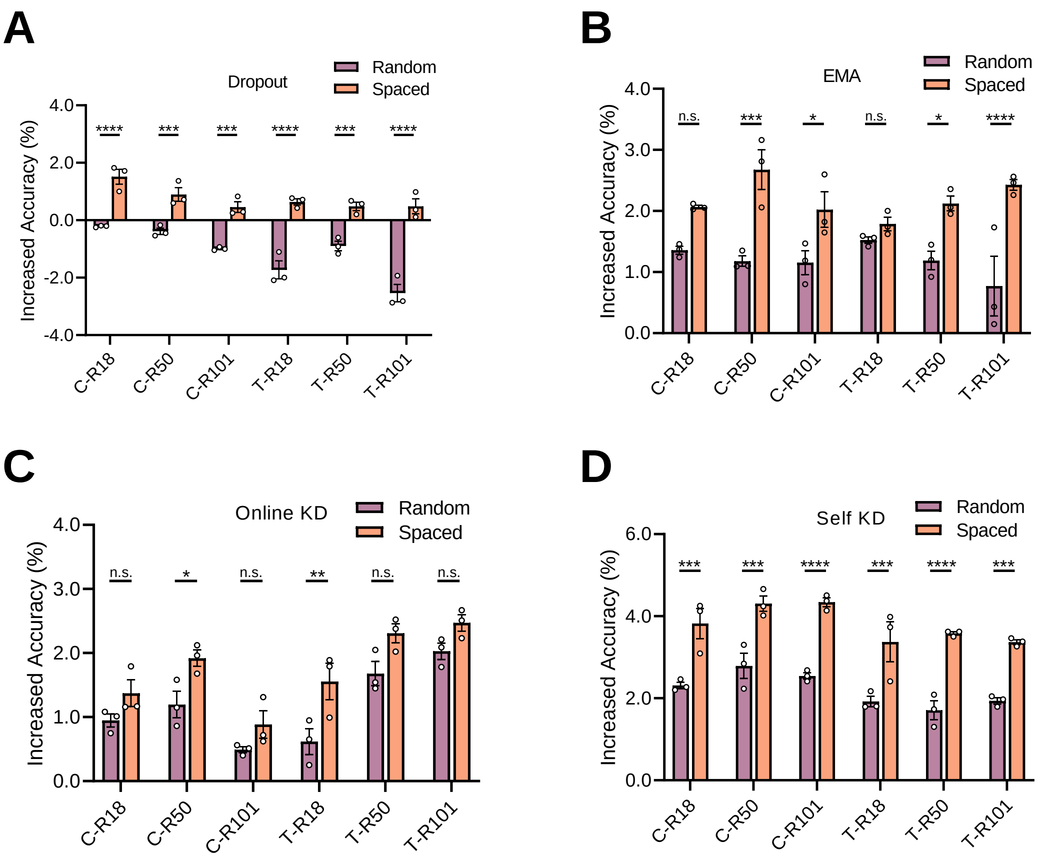


Figure S5: **Periodicity in spacing schedules is critical for enhanced generalization.** We compare the standard "Spaced" strategies (periodic intervals) against "Random" variants where the variation schedule was shuffled (irregular intervals) while maintaining the same average variation frequency. (**A**) Performance comparison for dropout. (**B**) Performance comparison for EMA. (**C**) Performance comparison for online KD. (**D**) Performance comparison for self KD. C, CIFAR-100. T, Tiny-ImageNet. R18, ResNet-18. R50, ResNet-50. R101, ResNet-101. All results are averaged over three runs with different random seeds. Data are presented as mean ± SEM. Statistical significance is determined by one-way ANOVA with Dunnett's multiple comparisons test. *p≤0.05, **p≤0.01, ****p≤0.0001, n.s., not significant.

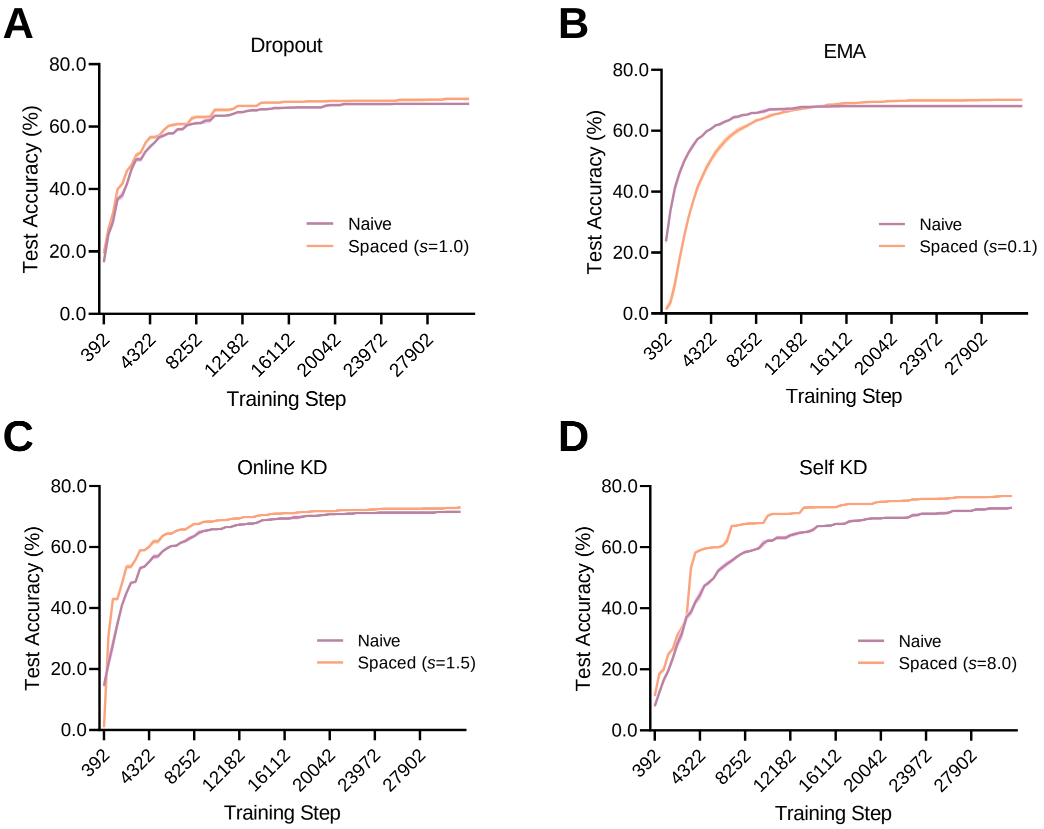


Figure S6: **Impact of spaced training on learning dynamics and performance ceiling.** We compare the test accuracy trajectories of standard "Naive" baselines versus their "Spaced" counterparts over training steps. (**A**) Training curve for dropout. (**B**) Training curve for EMA. (**C**) Training curve for online KD. (**D**) Training curve for self KD. All experiments use ResNet-18 on CIFAR-100. Data are averaged over three independent runs.

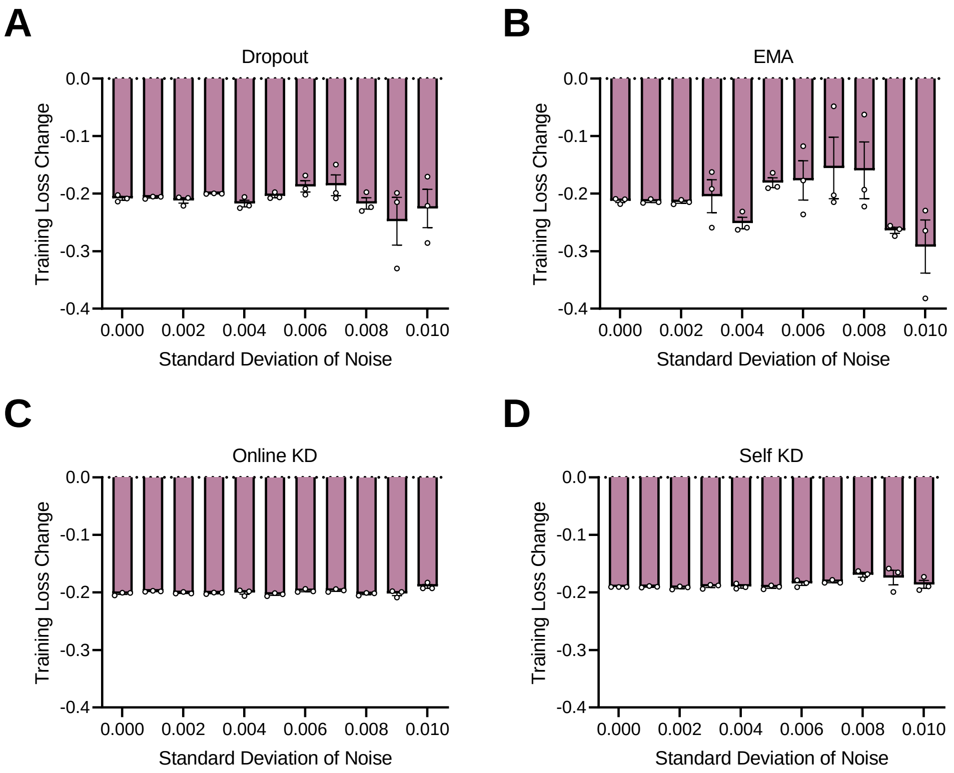


Figure S7: **Impact of parameter perturbations with Gaussian noise.** The bar display the training loss differences ($\Delta\mathcal{L} = \mathcal{L}_{\text{Spaced}} - \mathcal{L}_{\text{Naive}}$) between the standard "Naive" baselines and our "Spaced" counterparts. A negative value indicates that the Spaced variant experiences a smaller increase in training loss compared to the Naive baseline, suggesting a more robust parameterized solution. (**A**) Training loss differences for dropout. (**B**) Training loss differences for EMA. (**C**) Training loss differences for online KD. (**D**) Training loss differences for self KD. All experiments use ResNet-18 on CIFAR-100. Data are averaged over three independent runs.

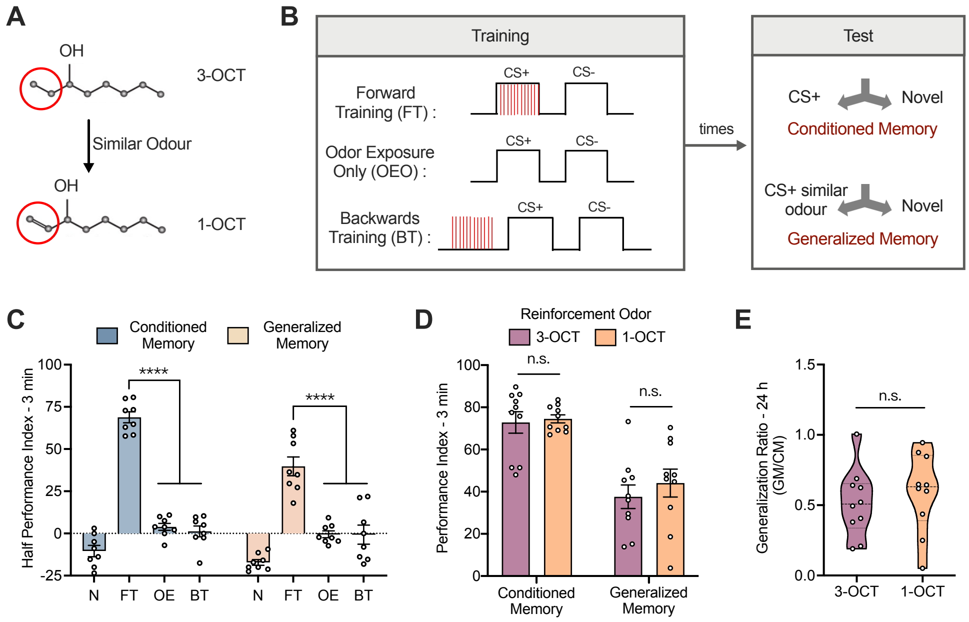


Figure S8: ***Drosophila* exhibit generalization to a similar odor after training.** (**A**) 3-OCT and 1-OCT are chemically similar odors. (**B**) Experimental paradigm for generalization training. Different groups of flies underwent forward training (FT), odor exposure only (OEO), or backwards training (BT). Conditioned and generalized memory are tested at indicated time points by presenting the CS+ odor (3-OCT) or a structurally similar odor (1-OCT), respectively, against a novel odor (MCH). (**C**) Both conditioned and generalized memory in the FT group are significantly higher than in control groups at 3 min. n=8. (**D**) Altering the reinforcement odor does not affect conditioned or generalized memory at 3 min. n=10. (**E**) Altering the reinforcement odor does not affect the generalization ratio at 3 min. n=10. Data are presented as mean ± SEM. Statistical significance is determined by two-way ANOVA with Sidak's multiple comparisons test or unpaired t test. ****$p \leq 0.0001$, n.s., not significant.

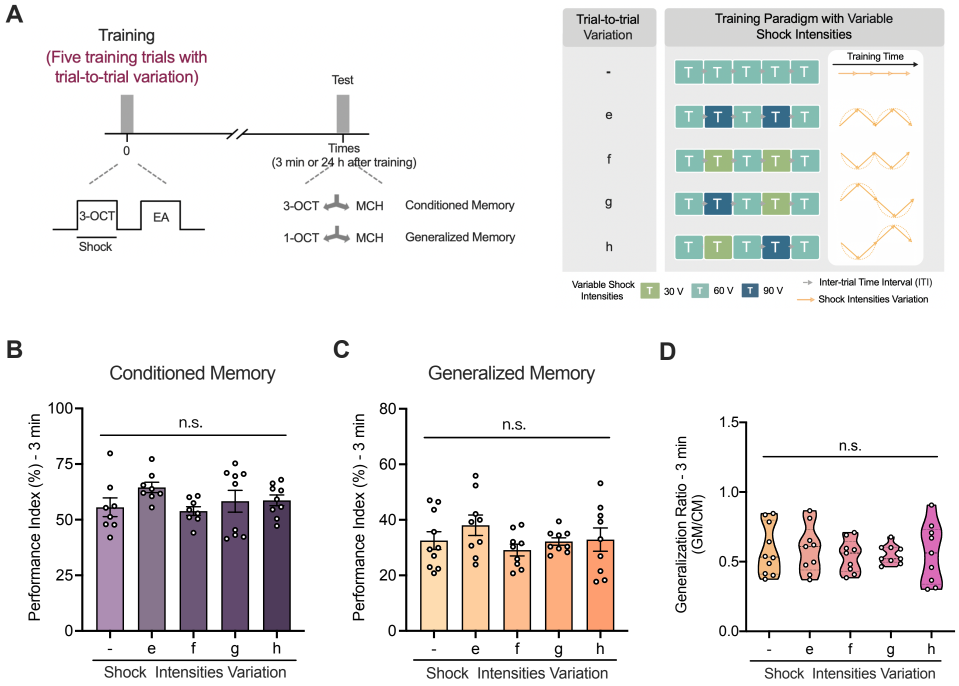


Figure S9: **Massed training with variable shock intensity has no effect on memory.** (**A**) Experimental paradigm for massed training with variable shock input. The control group (-) receives five trials with constant shock intensity, while experimental groups (e-h) receive trials with varying intensities to introduce encoding variability. (**B**-**D**) Conditioned memory (**B**, n=8-9), generalized memory (**C**, n=9-10), and generalization ratio (**D**, n=9-10) at 3 min are all no significant difference between control and experimental groups (e-h). Data are presented as mean ± SEM. Statistical significance is determined by one-way ANOVA with Dunnett's multiple comparisons test. n.s., not significant.

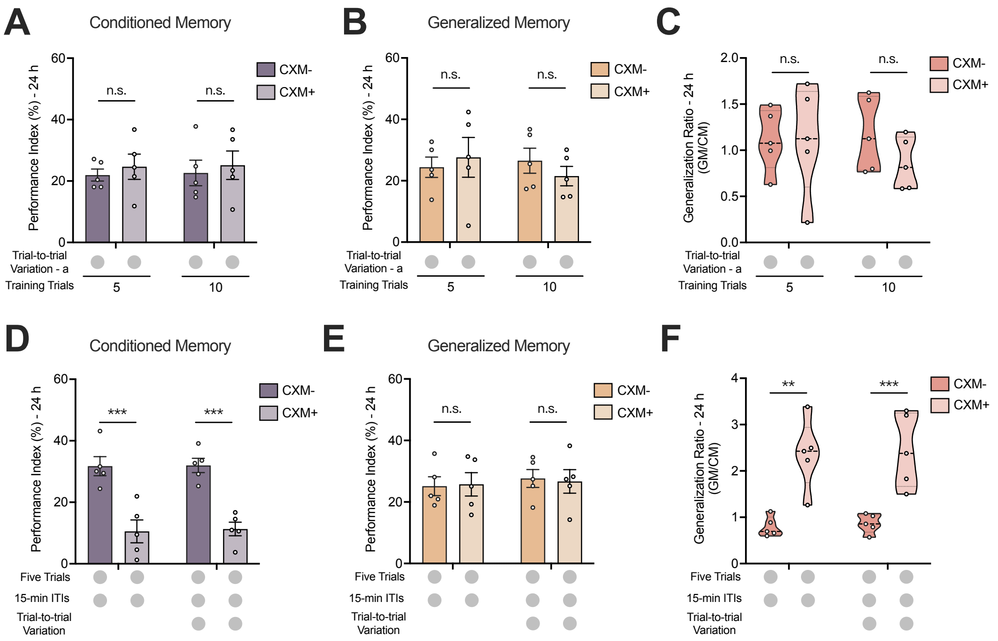


Figure S10: **Impact of protein synthesis inhibition on memory persistence and generalization.** (**A–C**) Trial-to-trial variation enhances generalization independently of protein synthesis during massed training. Conditioned memory (**A**, n=5), generalized memory (**B**, n=5), and generalization ratio (**C**, n=5) measured 24 h after 5 or 10 massed trials with trial-to-trial variation. No significant differences were observed between Control and CXM-treated groups. (**D–F**) Mechanistic dissociation between conditioned memory and generalization enhancement in spaced training. (**D**) CXM treatment significantly abolishes 24 h conditioned memory following spaced training compared to controls. n=5. (**E**) Generalized memory at 24 h remains intact in the CXM-treated group, showing no significant difference from controls. n=5. (**F**) The generalization ratio is significantly elevated in CXM-treated groups. n=5.

Table S1: **Detailed experimental results across multiple runs.** This table reports the absolute test accuracy (%) for three independent runs with different random seeds. We compare the "Naive" baselines against our proposed "Spaced" variants across four temporal ensemble strategies: Dropout (Fig. 2C, $s = 1.0$), EMA (Fig. 2G, $s = 0.1$), Online KD (Fig. 3C, $s = 1.5$), and Self KD (Fig. 3G, $s = 12.0$). Experiments were conducted on CIFAR-100 and Tiny-ImageNet datasets using ResNet-18, ResNet-50, and ResNet-101 backbones. The "Spaced" variants consistently achieve higher test accuracy compared to the naive baselines across all trials and network architectures.

| **Method** | **Dataset / Model** | **Naive** | | | **Spaced (Ours)** | | |
|---|---|---|---|---|---|---|---|
| | | Run 1 | Run 2 | Run 3 | Run 1 | Run 2 | Run 3 |
| Dropout | CIFAR-100 / ResNet-18 | 67.08 | 67.46 | 67.31 | 68.80 | 68.46 | 69.13 |
| | CIFAR-100 / ResNet-50 | 69.16 | 68.93 | 69.31 | 70.53 | 69.65 | 69.91 |
| | CIFAR-100 / ResNet-101 | 68.97 | 68.96 | 69.00 | 69.22 | 69.25 | 69.83 |
| | Tiny-ImageNet / ResNet-18 | 54.58 | 54.54 | 55.26 | 55.35 | 55.26 | 55.68 |
| | Tiny-ImageNet / ResNet-50 | 57.68 | 56.85 | 56.61 | 58.25 | 57.05 | 57.29 |
| | Tiny-ImageNet / ResNet-101 | 57.74 | 57.51 | 57.89 | 57.85 | 57.87 | 58.87 |
| EMA | CIFAR-100 / ResNet-18 | 68.32 | 68.17 | 67.87 | 70.35 | 70.29 | 69.92 |
| | CIFAR-100 / ResNet-50 | 68.81 | 68.72 | 69.12 | 71.97 | 71.53 | 71.18 |
| | CIFAR-100 / ResNet-101 | 67.62 | 67.44 | 67.53 | 70.22 | 69.26 | 69.18 |
| | Tiny-ImageNet / ResNet-18 | 61.63 | 61.74 | 61.56 | 63.63 | 63.48 | 63.18 |
| | Tiny-ImageNet / ResNet-50 | 64.10 | 64.49 | 64.30 | 66.46 | 66.55 | 66.25 |
| | Tiny-ImageNet / ResNet-101 | 63.95 | 64.13 | 64.10 | 66.21 | 66.69 | 66.56 |
| Online KD | CIFAR-100 / ResNet-18 | 70.97 | 70.99 | 71.20 | 72.76 | 72.16 | 72.36 |
| | CIFAR-100 / ResNet-50 | 71.77 | 71.72 | 72.05 | 73.89 | 73.68 | 73.73 |
| | CIFAR-100 / ResNet-101 | 71.70 | 72.35 | 72.03 | 73.01 | 73.07 | 72.65 |
| | Tiny-ImageNet / ResNet-18 | 59.23 | 59.50 | 58.85 | 61.02 | 60.49 | 60.74 |
| | Tiny-ImageNet / ResNet-50 | 61.11 | 61.23 | 60.64 | 63.49 | 63.25 | 63.16 |
| | Tiny-ImageNet / ResNet-101 | 61.52 | 61.27 | 61.09 | 63.75 | 63.78 | 63.76 |
| Self KD | CIFAR-100 / ResNet-18 | 72.98 | 72.80 | 74.09 | 77.10 | 77.05 | 77.18 |
| | CIFAR-100 / ResNet-50 | 75.60 | 75.20 | 75.73 | 79.84 | 79.86 | 79.75 |
| | CIFAR-100 / ResNet-101 | 76.34 | 76.14 | 75.93 | 80.48 | 80.51 | 80.44 |
| | Tiny-ImageNet / ResNet-18 | 61.51 | 60.62 | 61.10 | 63.92 | 64.35 | 65.08 |
| | Tiny-ImageNet / ResNet-50 | 63.79 | 63.48 | 63.58 | 66.77 | 67.08 | 67.08 |
| | Tiny-ImageNet / ResNet-101 | 62.99 | 63.90 | 63.47 | 67.49 | 67.49 | 67.09 |

**Algorithm 1** Spaced version of dropout

**Require**: model $f_\theta$, dataset $\mathcal{D}_{\text{train}}$, max dropout rate $p_{\max} \in [0, 1]$, epoch number $E$, space interval $s$

**Ensure**: train $f_\theta$ with periodically varying dropout rates

```
for 1 ≤ e ≤ E do
    Compute dropout rate p_e ← p_max · sin(2πe/s)
    for (x_i, y_i) ∈ D_train do
        Sample dropout mask r ∼ Bernoulli(p_e)
        Forward pass: h ← r ⊙ f_θ(x_i)
        Update θ ← θ − ∇_θ L_task(h, y_i)
    end for
end for
```

**Algorithm 2** Spaced version of EMA

**Require**: model $f_\theta$, dataset $\mathcal{D}_{\text{train}}$, momentum $\alpha$, epoch number $E$, space interval $s$

**Ensure**: train $f_\theta$ with momentum updates every $s$ steps

```
Initialize momentum model θ_m ← θ
Initialize step counter c ← 0
for 1 ≤ e ≤ E do
    for (x_i, y_i) ∈ D_train do
        Update θ ← θ − ∇_θ L_task(f_θ(x_i), y_i)
        c ← c + 1
        if c mod s = 0 then
            Update momentum model: θ_m ← αθ_m + (1 − α)θ
        end if
    end for
end for
```

**Algorithm 3** Spaced version of online KD

**Require**: student $f_\theta$, teacher $g_\phi$, dataset $\mathcal{D}_{\text{train}}$, distillation loss coefficient $\lambda$, epoch number $E$, space interval $s$

**Ensure**: train student model using spaced online KD

Initialize data index set: $\mathcal{R} \leftarrow \emptyset$
**for** $1 \leq e \leq E$ **do**
  **for** $(x_i, y_i) \in \mathcal{D}_{\text{train}}$ **do**
    $\mathcal{R} \leftarrow \mathcal{R} \cup \{i\}$
    Update teacher $\phi \leftarrow \phi - \nabla_\phi \mathcal{L}_{\text{task}}(g_\phi(x_i), y_i)$
    **if** $|\mathcal{R}| == s$ **then**
      **for** $j \in \mathcal{R}$ **do**
        Retrieve $(x_j, y_j)$ from $\mathcal{D}_{\text{train}}$
        Update student $\theta \leftarrow \theta - \nabla_\theta \left[\lambda \mathcal{L}_{\text{KD}}(f_\theta(x_j), g_\phi(x_j)) + (1-\lambda)\mathcal{L}_{\text{task}}(f_\theta(x_j), y_j)\right]$
      **end for**
      Clear index set: $\mathcal{R} \leftarrow \emptyset$
    **end if**
  **end for**
**end for**

**Algorithm 4** Spaced version of self KD

**Require**: model $f_\theta = f_{\theta_1} \circ \cdots \circ f_{\theta_m}$ with $m$ blocks, dataset $\mathcal{D}_{\text{train}}$, distillation loss coefficient $\lambda$, epoch number $E$, space interval $s$

**Ensure**: train $f_\theta$ using spaced self KD

Initialize data index set: $\mathcal{R} \leftarrow \emptyset$
**for** $1 \leq e \leq E$ **do**
  **for** $(x_i, y_i) \in \mathcal{D}_{\text{train}}$ **do**
    $\mathcal{R} \leftarrow \mathcal{R} \cup \{i\}$
    Update $\theta \leftarrow \theta - \nabla_\theta \mathcal{L}_{\text{task}}(f_\theta(x_i), y_i)$
    **if** $|\mathcal{R}| == s$ **then**
      **for** $j \in \mathcal{R}$ **do**
        Retrieve $(x_j, y_j)$ from $\mathcal{D}_{\text{train}}$
        Initialize $\mathcal{L}' \leftarrow \mathcal{L}_{\text{task}}(f_\theta(x_j), y_j)$
        **for** $1 \leq k < m$ **do**
          $\mathcal{L}' \leftarrow \mathcal{L}' + \lambda \mathcal{L}_{\text{KD}}(f_{\theta_1} \circ \cdots \circ f_{\theta_k}(x_j), f_\theta(x_j))$
        **end for**
        Update $\theta \leftarrow \theta - \nabla_\theta L'$
      **end for**
      Clear index set: $\mathcal{R} \leftarrow \emptyset$
    **end if**
  **end for**
**end for**
